# *Spd-2* Gene Duplication Suggests Cell-Type Specific Assembly Mechanisms of Pericentriolar Material

**DOI:** 10.1101/2021.10.27.466145

**Authors:** Ryan S. O’Neill, Afeez K. Sodeinde, Frances C. Welsh, Carey J. Fagerstrom, Brian J. Galletta, Nasser M. Rusan

**Author notes:** Contributed equally (Afeez K. Sodeinde, Frances C. Welsh). **Correspondence** (R.S.O.), (N.M.R.).

## Abstract

Centrosomes are multi-protein complexes that function as the major microtubule organizing center (MTOC) for the cell. While centrosomes play tissue-specific MTOC functions, little is known about how particular centrosome proteins are regulated across cell types to achieve these different functions. To investigate this cell type-specific diversity, we searched for gene duplications of centrosome genes in the *Drosophila* lineage with the aim of identifying centrosome gene duplications where each copy evolved for specialized functions. Through in depth functional analysis of a *Spd-2* gene duplication in the *Willistoni* group, we discovered differences in the regulation of PCM in somatic and male germline cells. The parental gene, *Spd-2A*, is expressed in somatic cells, where it can function to organize pericentriolar material (PCM) and the mitotic spindle in larval brain neuroblasts. Spd-2A is absent during male meiosis, and even when ectopically expressed in spermatocytes it fails to rescue PCM and spindle organization. In contrast, the new gene duplicate, *Spd-2B*, is expressed specifically in spermatocytes. During male meiosis, Spd-2B localizes to centrosomes, organizes PCM and spindles, and is sufficient for proper male fertility. Experiments using chimeric transgenes reveal that differences in the C-terminal tails of Spd-2A and Spd-2B are responsible for these functional changes. Thus, *Spd-2A* and *Spd-2B* have evolved complementary functions by specializing for distinct subsets of cells. Together, our results demonstrate that somatic cells and male germline cells have fundamentally different requirements for PCM, suggesting that PCM proteins such as Spd-2 is differentially regulated across cell types to satisfy distinct requirements.

**Highlights:**

- *Spd-2* gene duplication in the *Willistoni* group is rapidly evolving
- Parent gene *Spd-2A* is expressed in somatic cells, whereas the gene duplicate *Spd-2B* is expressed in spermatocytes
- Spd-2A organizes pericentriolar material during somatic cell mitosis
- Spd-2B is specialized for pericentriolar material organization during male meiosis

## Introduction

Centrosomes are complex organelles made up of many proteins, which together serve as the primary microtubule organizing centers (MTOCs) in most eukaryotic cells (Conduit et al., 2015; Vasquez-Limeta and Loncarek, 2021), ensuring proper spindle formation and chromosome segregation during cell division (Lerit and Poulton, 2016). The structure of the centrosome includes the core centrioles surrounded by a cloud of pericentriolar material (PCM), which anchors γ-Tubulin and nucleates microtubules (Conduit et al., 2015; Vasquez-Limeta and Loncarek, 2021). Although centrosomes function as MTOCs across many different cell types in an organism, the specific requirements for MTOC activity in a given cell type may be different as a result of cell type-specific characteristics such as cell size, cell cycle speed, or whether the cell undergoes mitotic or meiotic division (Greenan et al., 2010; Kwon and Scholey, 2004; Ohkura, 2015; Rieckhoff et al., 2019).

Relatively little is known about how centrosome proteins achieve cell type-specific differences in their functions. One way to accomplish cell type-specific functions is through alternative splicing. For example, in *Drosophila melanogaster* the centrosome gene *Cnn* encodes multiple splicing isoforms (Chen et al., 2017). Whereas most isoforms include a centrosome-targeting domain, one isoform instead includes a mitochondria-targeting domain which transforms the mitochondria into a MTOC during spermatogenesis. A second solution is through cell type-specific differences in binding partners. For example, in *D. melanogaster* PLP requires its binding partner calmodulin for its role at the basal body of neurons, while this interaction is dispensable for its role at the basal body of sperm (Galletta et al., 2014). Additionally, post-translational modifications such as phosphorylation or ubiquitination could alter centrosome protein function across cell types.

Another possible solution to modify centrosome protein functions is through gene duplication (Ohno, 1970; Taylor and Raes, 2004). For a single copy gene, mutations that enhances protein function in one cell type may have a detrimental effect on its function in other cell types, thus limiting the potential for single copy genes to achieve optimal function across all cell types. Gene duplication can bypass this limitation by allowing each duplicate to accumulate mutations independently and become specialized for a subset of functions (Des Marais and Rausher, 2008; Hughes, 1994). Some centrosome genes appear to have evolved in this way. For example, a gene duplication of CEP135 in vertebrates gave rise to TSGA10, which has testis-specific expression and appears to be critical for the formation of sperm head-tail attachment (Modarressi et al., 2001; Sha et al., 2018). We reasoned that by identifying and studying centrosome gene duplications that evolved towards cell type-specific specialization, we could gain insight into the nature of cell type-specific differences in MTOC activity, how centrosome protein functions change across cell types, and the mechanisms underlying cell type-specific centrosome protein regulation.

Here, we present our work on a duplication of the *Spd-2* gene in *Drosophila willistoni*. In *Caenorhabditis elegans* and humans, SPD-2/CEP192 are required for both PCM recruitment and centriole duplication (Gomez-Ferreria and Sharp, 2008; Kemp et al., 2004; Pelletier et al., 2004), whereas *D. melanogaster Spd-2* is required for PCM recruitment but not for centriole duplication (Dix and Raff, 2007; Giansanti et al., 2008). Spd-2 localizes to the centrosomes where it is required for proper PCM organization in both larval brain neuroblast mitosis and male spermatocyte meiosis (Dix and Raff, 2007; Giansanti et al., 2008). Spd-2 is initially recruited to the centriole wall by the bridge zone protein Asl (Conduit et al., 2014). Once recruited to the centrosome, Spd-2 in turn recruits another PCM protein Cnn, and together they organize a stable PCM for γ-Tubulin recruitment and microtubule nucleation (Conduit et al., 2010; Conduit et al., 2014; Goshima et al., 2007). However, many questions remain as to how Spd-2 is regulated, and it is unknown whether Spd-2 functions differently in the contexts of mitosis versus meiosis.

Our investigation of *D. willistoni Spd-2* gene duplicates reveals that the parental gene, *Spd-2A*, maintained its ability to organize PCM during mitosis but lost the ability to organize PCM during meiosis. In contrast, the rapidly evolving new gene copy, *Spd-2B*, is specifically expressed in the testes where it functions to organize meiotic PCM. Experiments using chimeric transgenes reveal that the amino acid changes responsible for differences in meiotic PCM assembly function occurred in the C-terminal tail of the Spd-2 duplicates, suggesting that the C-terminal region of Spd-2 is a target of regulation to achieve differences in PCM function between mitotic and meiotic cell division.

## Results

### Screen for Centrosome Gene Duplications in the *Drosophila* Lineage

To identify duplications of centrosome genes in the *Drosophila* lineage, we used BLAST to screen 13 *D. melanogaster* centrosome proteins (Figure 1A) against the annotated protein sequence databases of 35 *Drosophila* species (Supplemental File 1). Of the 13 genes, five have undergone duplication at least once (Figure 1B). *Sas-4*, which encodes a core centriolar protein, has a nearly identical tandem duplication in *Drosophila yakuba*. *Asl*, *Cnn*, and *Cp110* were all duplicated multiple times. Interestingly, both *Asl* duplications (Asl-B and Asl-C) were subsequently re-duplicated in different lineages (Figure 1B, Asl). The *Cnn* duplication in *Drosophila miranda* was nearly identical to the parent gene, whereas a second duplication found in several species was a highly divergent partial duplication containing only a small portion of the N-terminal protein coding sequence (Figure 1B, Cnn). Finally, *Spd-2* was duplicated in both *D. melanogaster* and *D. willistoni*. In *D. melanogaster*, we found the previously identified chimeric pseudogene *CR18217*, which contains an N-terminal fragment of *Spd-2* along with the NUDIX domain from *CG4098* (Rogers and Hartl, 2012). In *D. willistoni*, the new copy of *Spd-2* encoded a nearly full length protein that showed substantial divergence from its parent gene. We chose the *Spd-2* gene duplication for detailed analysis, and herein refer to the *D. willistoni* parent gene as *Spd-2A*, the *D. willistoni* new gene duplicate as *Spd-2B*, and the *D. melanogaster* single copy gene as *Spd-2*.

**Figure 1.**
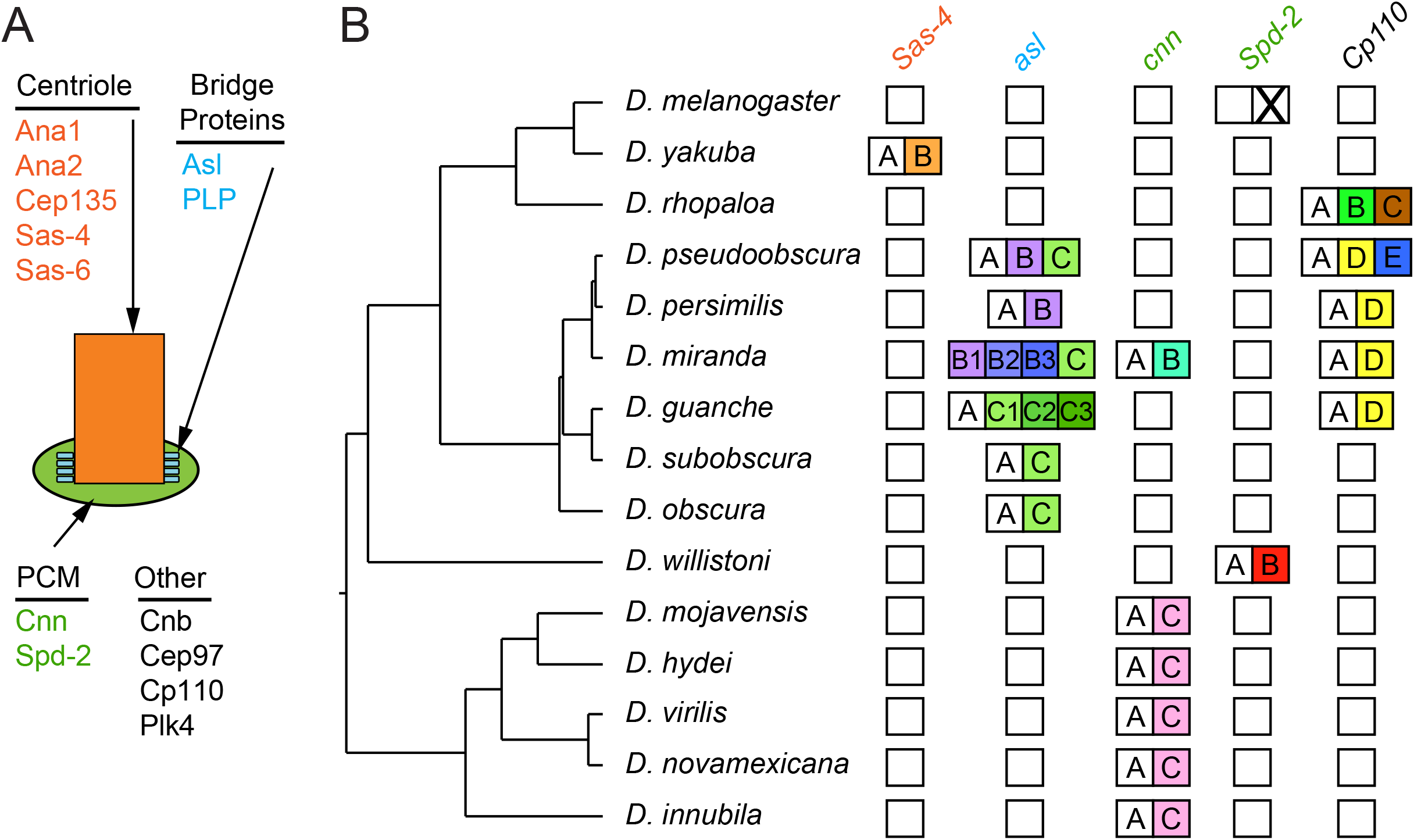
Centrosome gene duplications in the *Drosophila* lineage. (A) Diagram of a centrosome showing proteins used in a BLAST screen for centrosome gene duplications. Categories included core centriole proteins (orange), bridge proteins (blue), and PCM proteins (green). (B) Summary of BLAST screen results. Only genes and species where duplications were found are shown. White boxes represent single copy genes. Lettered boxes represent duplicated genes: a white box with “A” represents the orthologs of the single copy gene, while each colored boxes with a letter represents a unique duplication. For *Spd-2*, duplication “X” in *D. melanogaster* represents a chimeric pseudogene. For *asl*, duplications “B” and “C” were subsequently re-duplicated multiple times, and these duplications are represented by boxes with a letter and number.

### *Spd-2B* is a Rapidly Evolving Duplication of *Spd-2A*

We obtained the genome sequence upstream and downstream of *Spd-2A* and used BLAST to determine the corresponding regions in the *D. melanogaster* genome, revealing the same upstream and downstream genes for both *Spd-2A* and *Spd-2* and thus confirming their orthology (Figure 2A). We used the same strategy to identify the locus of *Spd-2B*, which was between the *bug* and *Mlf* genes in *D. willistoni*; in *D. melanogaster*, *bug* and *Mlf* are directly adjacent on Chromosome 2 (Figure 2A).

**Figure 2.**
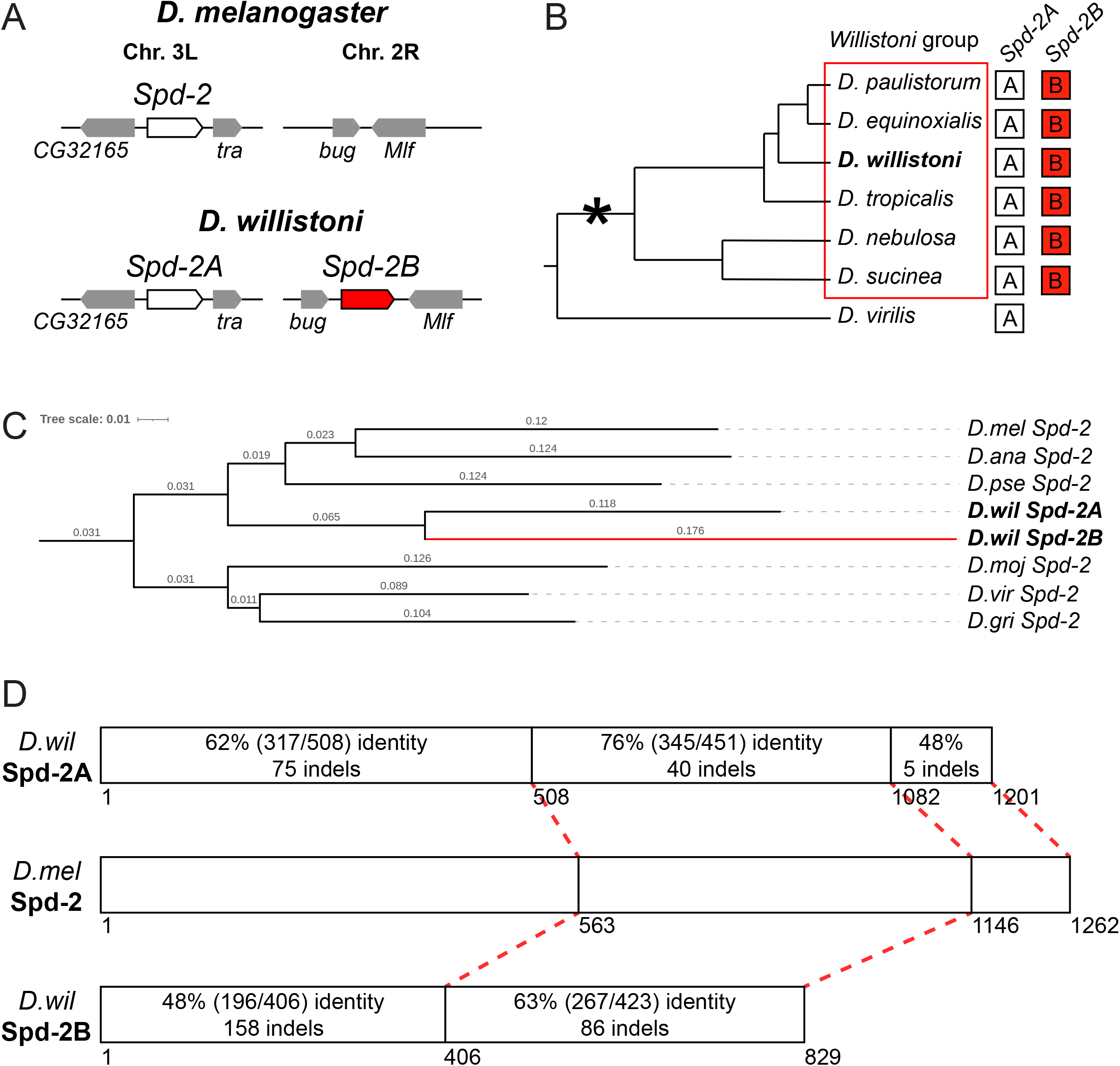
*Drosophila willistoni Spd-2B* is a conserved, rapidly evolving duplication of *Spd-2*. (A) Top: *D. melanogaster* gene regions of *Spd-2* locus (Chromosome 3L, left) and the syntenic locus to where *Spd-2B* is inserted in *D. willistoni* (Chromosome 2R, right). Bottom: *D. willistoni* gene regions of *Spd-2A* (left) and *Spd-2B* (right). Adapted from Flybase. (B) *Willistoni* group lineage showing presence of both *Spd-2A* and *Spd-2B* in multiple species confirmed by PCR test. Note the absence of *Spd-2B* in the outgroup species *Drosophila virilis*. (C) Phylogenetic tree based on a trimmed codon alignment of *Spd-2*, *Spd-2A* and *Spd-2B*. The branch length of *Spd-2B* (red branch) is significantly longer than *Spd-2A*, indicating rapid divergence. (D) Diagrammatic representation of *D. melanogaster* Spd-2 (middle) compared to *D. willistoni* Spd-2A (top) and Spd-2B (bottom) proteins. A comparison of amino acid identity and indels to *D. melanogaster* Spd-2 are shown for both Spd-2A and Spd-2B. Spd-2B is completely lacking the Spd-2 C-terminal tail.

To determine whether *Spd-2B* is a *D. willistoni*-specific duplication we used PCR and sequencing to screen for *Spd-2A* and *Spd-2B* in six species of the *Willistoni* group. PCR screening revealed that both *Spd-2A* and *Spd-2B* were present in all six species (Figure 2B, Figure S1A), whereas *Spd-2B* was not found in the outgroup species *D. virilis*. Thus, the duplication that gave rise to *Spd-2B* likely occurred in the ancestor of the *Willistoni* group, between ∼22-50 million years ago (Russo et al., 2013), and was subsequently retained in all daughter species.

We next assessed the sequence conservation of *Spd-2A* compared to *Spd-2B*. We generated a codon alignment of *Spd-2* homologs from seven species. This codon alignment revealed that *Spd-2B* is evolving at a significantly faster rate compared to Spd-2A (Figure 2C). To assess conservation of protein sequence and structure, we compare Spd-2A and Spd-2B to Spd-2 protein sequences (Figure 2D). Compared to Spd2, Spd-2B is less conserved than Spd-2A and contains over twice the number of amino acid indels. The C-terminal halves of Spd-2A and Spd-2B are more strongly conserved compared to their N-terminal halves. However, Spd-2B is missing the C-terminal ∼119 amino acid tail of Spd-2A and Spd-2. Together, these results indicate that *Spd-2B* is evolving more rapidly than *Spd-2A*, suggesting functional divergence or specialization for *Spd-2B*.

### Spd-2A, but not Spd-2B, organizes PCM and the Mitotic Spindle in Larval Brain Neuroblasts

To explore the functions of Spd-2A and Spd-2B, we generated *D. melanogaster* animals expressing GFP-tagged transgenes of *Spd-2A*, *Spd-2B*, and *Spd-2*. To preserve major regulatory elements and reflect native expression patterns, each transgene included their complete native upstream and downstream intergenic DNA sequences and were inserted into the same insulated attP landing site in the *D. melanogaster* genome (Figure S2A). Whereas both native promoter *mel(p)-*Spd-2::GFP and *wil(p)-*Spd-2A::GFP were expressed in the larval brain neuroblasts (Figure 3A, B), *wil(p)-*Spd-2B::GFP was not detectable (Figure 3C). Similarly, *mel(p)-*Spd-2::GFP and *wil(p)-*Spd-2A::GFP, but not *wil(p)-*Spd-2B::GFP, were expressed in larval wing discs (Figure S2B-D). Thus, Spd-2 and Spd-2A are expressed in somatic tissues, whereas Spd-2B is absent.

**Figure 3.**
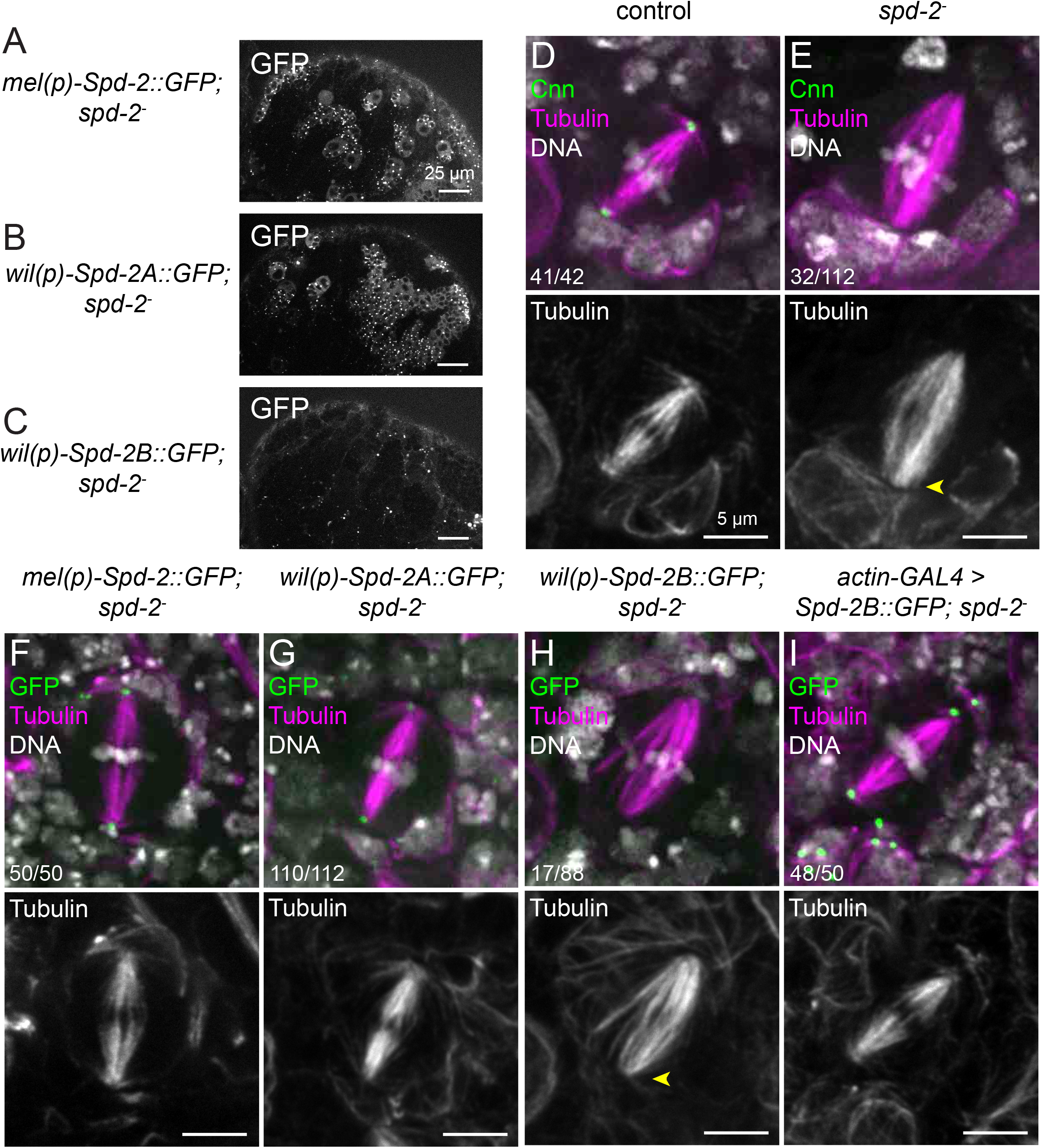
Spd-2A and Spd-2B can organize the mitotic spindle in larval brain neuroblasts. (A-C) Micrographs of unfixed 3^rd^ instar larval brains expressing *D. melanogaster* Spd-2::GFP (A) and *D. willistoni* Spd-2A::GFP (B) or Spd-2B::GFP (C). (D-E) Micrographs of neuroblast metaphase spindles of controls (D) and *spd-2^-^* mutants (E), stained for Cnn (green), Tubulin (magenta), and DNA (white). Single channel micrographs of Tubulin alone are shown below. Controls show normal morphology with a tight spindle that is properly focused at the poles (1/42 unfocused). *spd-2^-^* mutants have a barrel shaped spindle with frequently unfocused poles (32/112 unfocused; yellow arrowhead). (F-I) Micrographs of neuroblast metaphase spindles from *spd-2^-^* mutants with rescue constructs *mel(p)-Spd-2::GFP* (F), *wil(p)-Spd-2A::GFP* (G), *wil(p)-Spd-2B::GFP* (H), and *actin-GAL4 > UAS-Spd-2B::GFP* (I) stained for Tubulin (magenta), and DNA (white), and expressing Spd-2(A/B)::GFP (green). Both Spd-2::GFP and Spd-2A::GFP are expressed in neuroblasts and rescue *spd-2^-^* spindle morphology (0/50 and 2/112 unfocused, respectively; F, G). Spd-2B::GFP is not detectable in neuroblasts, and fails to rescue *spd-2^-^* spindle morphology (17/88 unfocused; H, yellow arrowhead). When expressed via *actin-GAL4*, Spd-2B::GFP localizes to the poles and properly organizes the spindle (2/50 unfocused; I).

Larval brain neuroblasts in metaphase have well organized mitotic spindle poles and radial astral microtubules (Figure 3D). Similar to previous reports, we found that *spd-2^-^* neuroblast metaphase spindles were abnormally barrel shaped with unfocused poles that also lack astral microtubules (Figure 3E; Giansanti et al., 2008). Expressing *mel(p)-Spd-2::GFP* or *wil(p)-Spd-2A::GFP* transgenes in the *spd-2^-^* mutant background rescued these abnormal metaphase spindle defects (Figure 3F, G). However, *wil(p)-Spd-2B::GFP* transgene failed to rescue the *spd-2^-^* spindle phenotypes (Figure 3H), consistent with the lack of GFP expression in larval brain neuroblasts. Ectopically expressing a *UAS-Spd-2B::GFP* transgene using *actin-GAL4* in the *spd-2^-^* background fully rescued the mutant spindle defects (Figure 3I). This indicates that Spd-2B protein is capable of performing Spd-2 or Spd-2A function in the brain, but *Spd-2B* is transcriptionally inactive in the brain.

We next wondered if Spd-2A could fully rescue centrosome assembly in somatic cells. During neuroblast mitosis, Spd-2 is known to recruit and stabilize Cnn to organize and expand the PCM (Conduit et al., 2010; Conduit et al., 2014; Goshima et al., 2007). We found that, similar to previous reports, *spd-2^-^* neuroblasts are often missing centrosomes and fail to robustly recruit γ-Tubulin and Cnn as compared to controls (Figure 4A, B, G-I; Conduit et al., 2007; Giansanti et al., 2008; Goshima et al., 2007). Both *mel(p)-*Spd-2::GFP and *wil(p)-*Spd-2A::GFP localize to centrosomes and rescue *spd-2^-^* recruitment of both γ-Tubulin and Cnn (Figure 4C, D). As expected, *wil(p)-*Spd-2B::GFP is undetectable in larval brain neuroblasts, and fails to rescue *spd-2^-^* centrosome numbers, γ-Tubulin recruitment or Cnn recruitment (Figure 4E). However, consistent with mitotic spindle rescue, *UAS-*Spd-2B::GFP ectopic expression via *actin-GAL4* localizes to centrosomes and fully rescues *spd-2^-^* centrosome number and recruitment of γ-Tubulin and Cnn (Figure 4F-I). Together these results show that, although Spd-2B is not normally expressed in larval brain cells, the encoded protein is fully capable of organizing the mitotic spindle and PCM via Cnn and γ-Tubulin recruitment.

**Figure 4.**
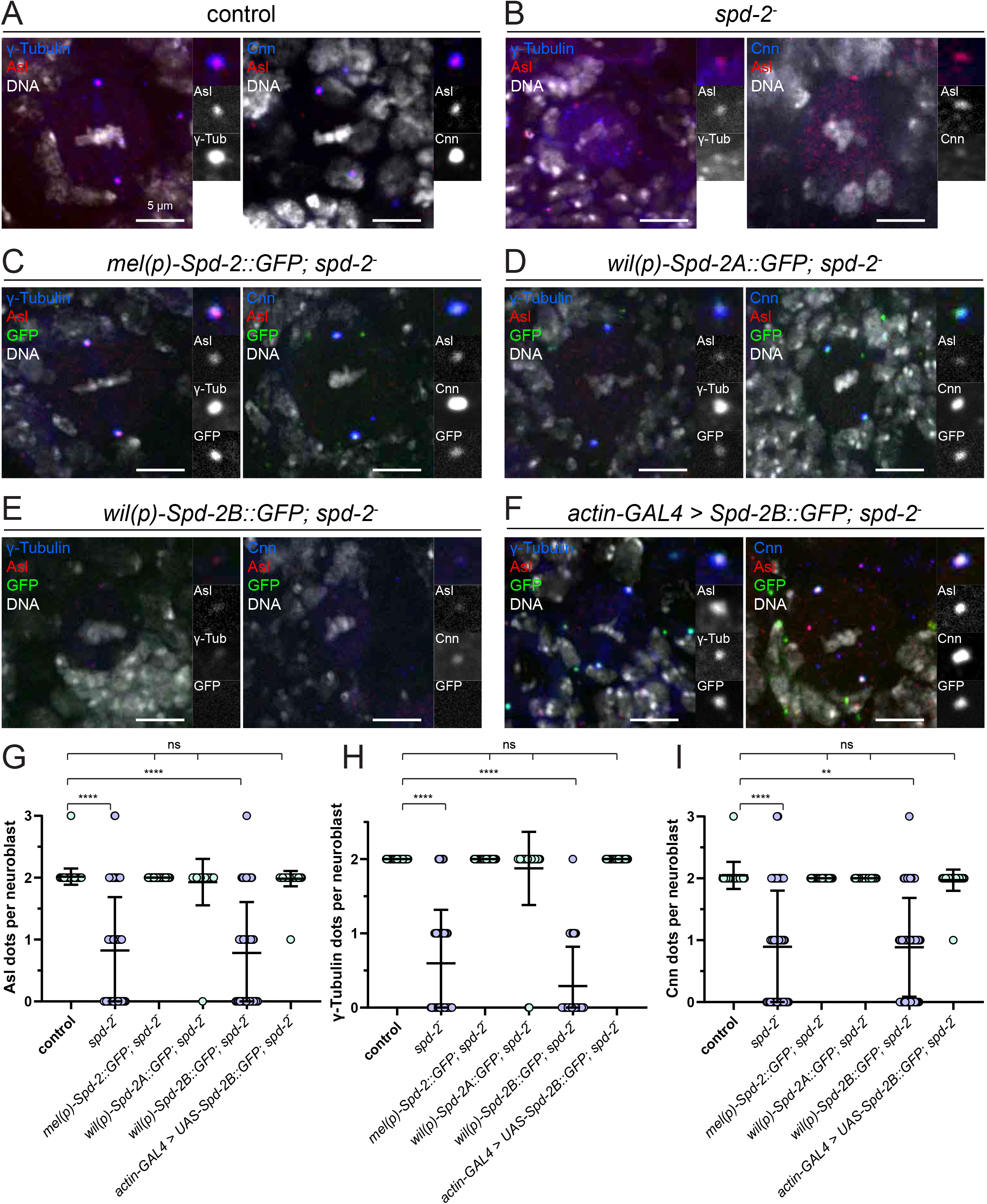
Spd-2A and Spd-2B can recruit centrosomal γ-Tubulin and Cnn in larval brain neuroblasts. (A-B) Micrographs of metaphase neuroblasts from controls (A) and *spd-2^-^* mutants mutants (B), stained for Asl (red), DNA (white), and either γ-Tubulin (blue, left panels) or Cnn (blue, right panels). A zoom to the apical centrosome with isolated Asl and γ-Tubulin or Cnn channels are shown to the right. (C-F) Micrographs of metaphase neuroblasts from *spd-2^-^* mutants mutants with rescue constructs *mel(p)-Spd-2::GFP* (C), *wil(p)-Spd-2A::GFP* (D), *wil(p)-Spd-2B::GFP* (E), or *actin-GAL4 > UAS-Spd-2B::GFP* (F) expressing Spd-2(A/B)::GFP (green) and stained Asl (red), DNA (white), and either γ-Tubulin (blue, left panels) or Cnn (blue, right panels). (G-I) Quantification of the number of Asl dots (G), γ-Tubulin dots (H), and Cnn dots (I) per neuroblast for genotypes in A-F. Error bars represent standard deviation. Significance tested via ordinary one-way ANOVA. ns = not significant; **** p < 0.001; ** p < 0.01.

### Spd-2A and Spd-2B have complementary centriolar localizations during spermatogenesis

In addition to its roles in larval brain neuroblasts, Spd-2 has also been studied for its role in male meiosis. First, we characterized the expression patterns of native promoter transgenes by looking at centrosomal localization of Spd-2::GFP at various stages during spermatogenesis. These stages included the germline stem cells and spermatogonia, which undergo mitotic divisions (Figure 5A), mature pre-meiotic spermatocytes (Figure 5B), and spermatocytes, which undergo meiosis (Figure 5C). During mitotic stages, both *mel(p)-*Spd-2::GFP and *wil(p)-*Spd-2A::GFP localized to centrosomes and expanded into PCM, whereas *wil(p)-*Spd-2B::GFP was barely detectable (Figure 5A). In pre-meiotic mature spermatocytes this pattern is reversed, with *mel(p)-*Spd-2::GFP and *wil(p)-*Spd-2B::GFP localized to centrosomes, whereas *wil(p)-*Spd-2A::GFP was now barely detectable (Figure 5B). Finally, this pattern continued through the meiotic divisions, with *mel(p)-*Spd-2::GFP and *wil(p)-*Spd-2B::GFP localized to centrosomes and expanding into PCM, and *wil(p)-*Spd-2A::GFP was still barely detectable (Figure 5C). The cytoplasmic level of Spd-2A::GFP was equal in mitotic and meiotic stages (Figure 5D), indicating that the reduction in centrosomal Spd-2A::GFP was not due to total cellular protein degradation but rather a delocalization of Spd-2A::GFP off of the centrosome. The increase of Spd-2B::GFP on the centrosome and PCM in later stages was accompanied by a global cytoplasmic increase of Spd-2B::GFP (Figure 5D), suggesting transcriptional or translational control of Spd-2B::GFP. Nonetheless, the levels of cytoplasmic Spd-2B::GFP were no greater than Spd-2A::GFP in meiotic spermatocytes, yet only Spd-2B::GFP is loaded onto the centriole and expanded into the PCM, strongly suggesting that Spd2A and Spd2B protein are differentially regulated, possibly via unique phosphorylation sites. Together, these results show that Spd-2A and Spd-2B have complementary patterns of centrosomal localization such that a combination of both are required to recapitulate the full localization of Spd-2 throughout all stages of spermatogenesis.

**Figure 5.**
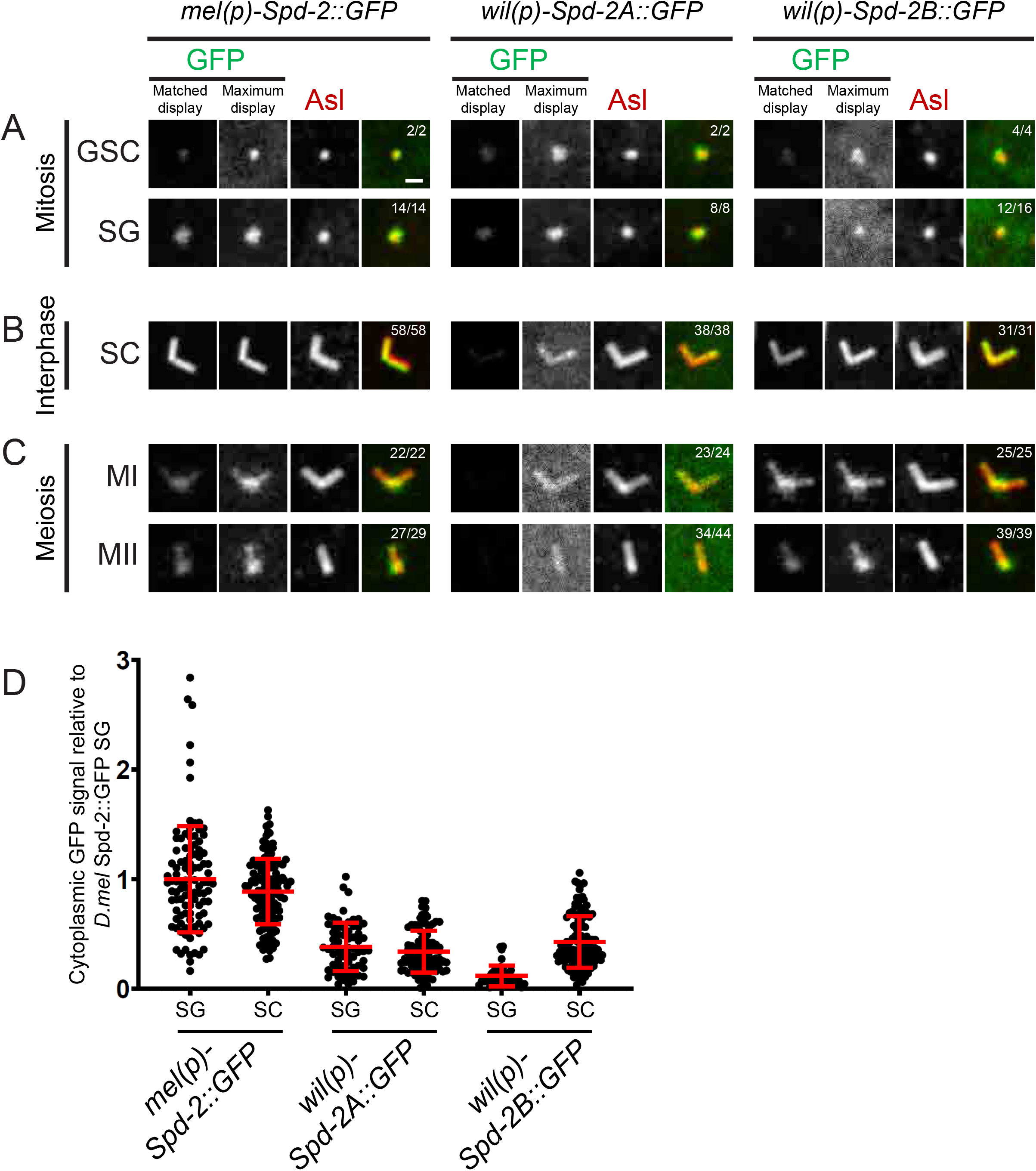
Spd-2A and Spd-2B have complementary centriolar localizations in spermatogenesis. (A, B, C) Micrographs of *spd-2^-^* mutants with *mel(p)-Spd-2::GFP*, *wil(p)-Spd-2A::GFP*, or *wil(p)-Spd-2B::GFP* (GFP in green) stained for Asl (red) in various stages of spermatogenesis: (A) mitotic stages of male germline stem cells (GSC) and spermatogonia (SG); (B) interphase stage of mature spermatocytes (SC); and (C) spermatocytes in meiosis I (MI); meiosis II (MII). GFP channels are displayed in two ways: first, using the same display range for all images (Matched display, left), and, second, such that the dynamic range of each image corresponds to the brightest and dimmest pixels (Maximum display, right). Note that Spd-2::GFP has robust centrosomal localization throughout spermatogenesis (A). In constrast, Spd-2A::GFP localizes to the centrosome early (B), whereas Spd-2B::GFP localizes to the centrosome late in spermatogenesis (C). (D) Quantification of cytoplasmic Spd-2(A/B)::GFP in spermatogonia (SG) and spermatocytes (SC). All values are normalized to *D. melanogaster* Spd-2 in SG. Note that Spd-2A cytoplasmic levels remain relatively consistent throughout spermatogenesis, while centrosomal levels appear reduced during meiosis.

### Spd-2B is Specialized for PCM Organization During Male Meiosis

Next, we investigated the PCM recruitment functions of Spd-2A and Spd-2B. Wild type spermatocytes recruit γ-Tubulin into the PCM during meiosis I (Figure 6A), whereas *spd-2^-^* spermatocytes only recruit a small amount of γ-Tubulin that fails to spread into PCM (Figure 6B). *mel(p)-Spd-2::GFP* rescues *spd-2^-^* γ- Tubulin recruitment, causing a similar sized PCM, albeit with higher than wild type γ-Tubulin levels (Figure 6C). As expected based on lack of localization, *wil(p)-Spd-2A::GFP* fails to rescue *spd-2^-^* γ-Tubulin recruitment (Figure 6D). In contrast, *wil(p)-Spd-2B::GFP* rescues *spd-2^-^* γ-Tubulin recruitment, causing a similar sized centrosome with similar γ-Tubulin levels compared to controls, but qualitatively less than *mel(p)-Spd-2::GFP* (Figure 6E). We also investigated the meiotic spindle organizing functions of Spd-2A and Spd-2B. In meiosis I, controls organize a proper meiotic spindle with robust asters (Figure 6F), whereas *spd-2^-^* spermatocyte centrosomes are abnormal, recruiting very weak or no astral microtubules, and are frequently detached from the poles (Figure 6G, H). *mel(p)-Spd-2::GFP* fully rescues *spd-2^-^* meiotic spindle defects (Figure 6I). As expected, *wil(p)-Spd-2A::GFP* fails to rescue *spd-2^-^* meiotic spindle defects (Figure 6J) whereas *wil(p)-Spd-2B::GFP* fully rescues (Figure 6K). Thus, consistent with localization patterns (Figure 5), Spd-2B, but not Spd2A, controls PCM assembly and spindle organizing functions during male meiosis.

**Figure 6.**
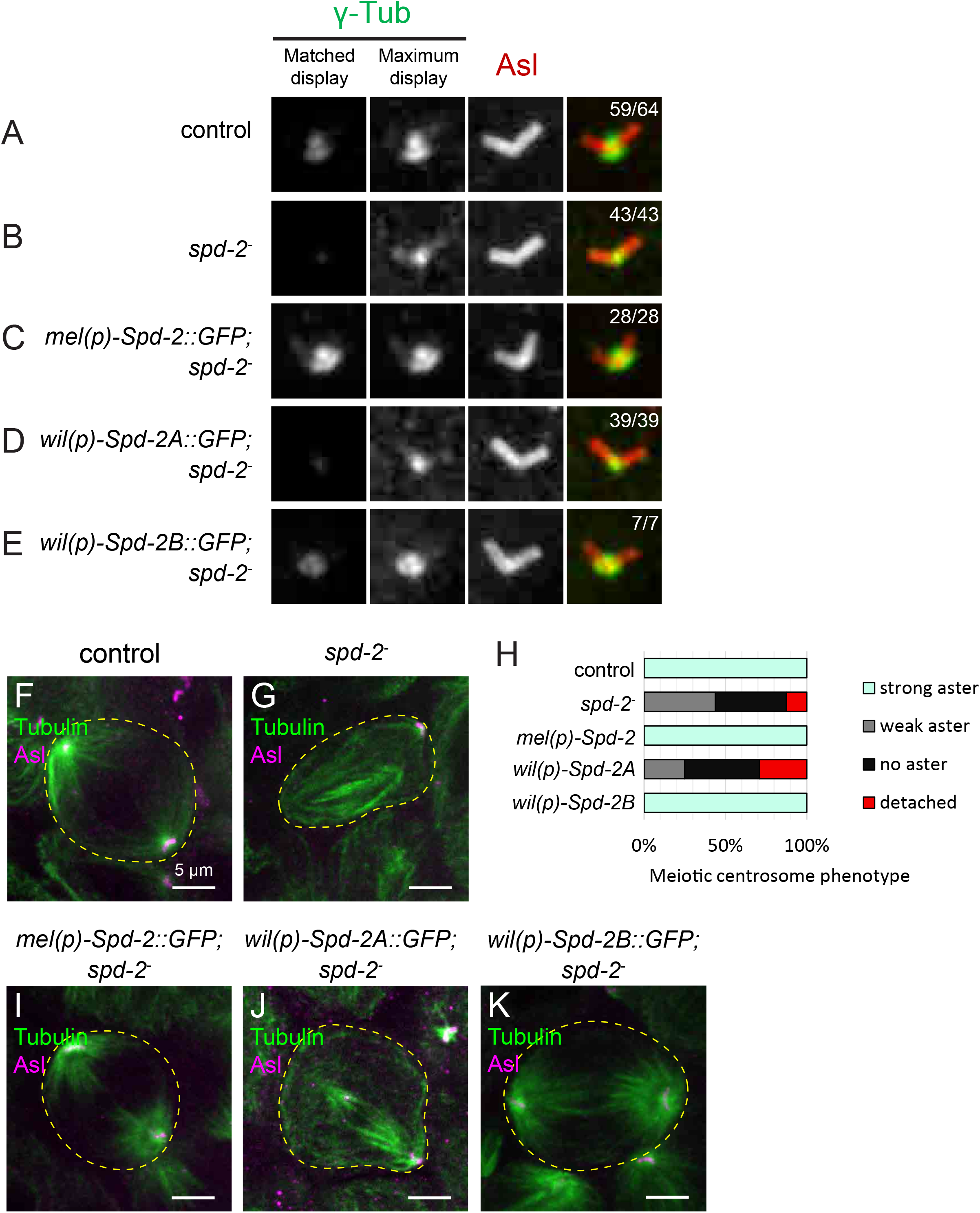
Spd-2B, but not Spd-2A, recruits γ-Tubulin and organizes spindles during meiosis. (A-E) Micrographs of control (A), *spd-2^-^* mutant (B), and *mel(p)-Spd-2::GFP; spd-2^-^* (C) *wil(p)-Spd-2A::GFP; spd-2^-^* (D) and *wil(p)-Spd-2B::GFP; spd-2^-^* (E) spermatocyte centrosomes during meiosis I, stained for γ-Tubulin (green) and Asl (red). γ-tubulin channels are displayed in two ways: first, using the same display range for all images (Matched display, left), and, second, such that the dynamic range of each image corresponds to the brightest and dimmest pixels (Maximum display, right). Note that controls (A) and Spd-2B::GFP rescues (E) recruit similar levels of γ-Tubulin to the centrosomes, whereas both Spd-2 mutants (B) and Spd-2A::GFP (D) rescues fail to properly recruit γ-Tubulin. (F, G) Micrographs of control (F) and *spd-2^-^* mutant (G) spermatocytes during meiosis I, stained for Tubulin (green) and Asl (magenta). (H) Quantification of centrosome phenotypes. Strong aster corresponds to centrosomes with full asters as seen in (F); weak asters correspond to centrosomes with only a few microtubules as seen in (G); no asters corresponds to centrosomes with no visible microtubules; and detached corresponds to centrosomes that have detached from the poles. (I-K) Micrographs of *spd-2^-^* mutants with native promoter rescues *mel(p)-Spd-2::GFP* (I), *wil(p)-Spd-2A::GFP* (J), or *wil(p)-Spd-2B::GFP* (K) stained for Tubulin (green) and Asl (magenta).

To further test the functions of Spd-2A and Spd-2B in spermatogenesis, we tested whether they were sufficient to rescue *spd-2^-^* male sterility. We allowed individual males to mate with two virgin females over the course of four days, and then counted the total number of offspring (Figure 7A). Wild type males produced an average 127 offspring. As previously reported, *spd-2^-^* males were completely sterile, producing zero offspring (Dix and Raff, 2007; Giansanti et al., 2008). *mel(p)-Spd-2::GFP* rescued *spd-2^-^* male sterility, with an average 95 offspring per male. As expected based on localization and ability to rescue γ-Tubulin recruitment, *wil(p)-Spd-2A::GFP* completely failed to rescue *spd-2^-^* male sterility. In contrast, *wil(p)-Spd-2B::GFP* rescued *spd-2^-^* male sterility similar to *mel(p)-Spd-2::GFP*, with an average 99 offspring per male. Together, along with γ-Tubulin rescue these results show that *Spd-2B* is specialized for spermatogenesis.

**Figure 7.**
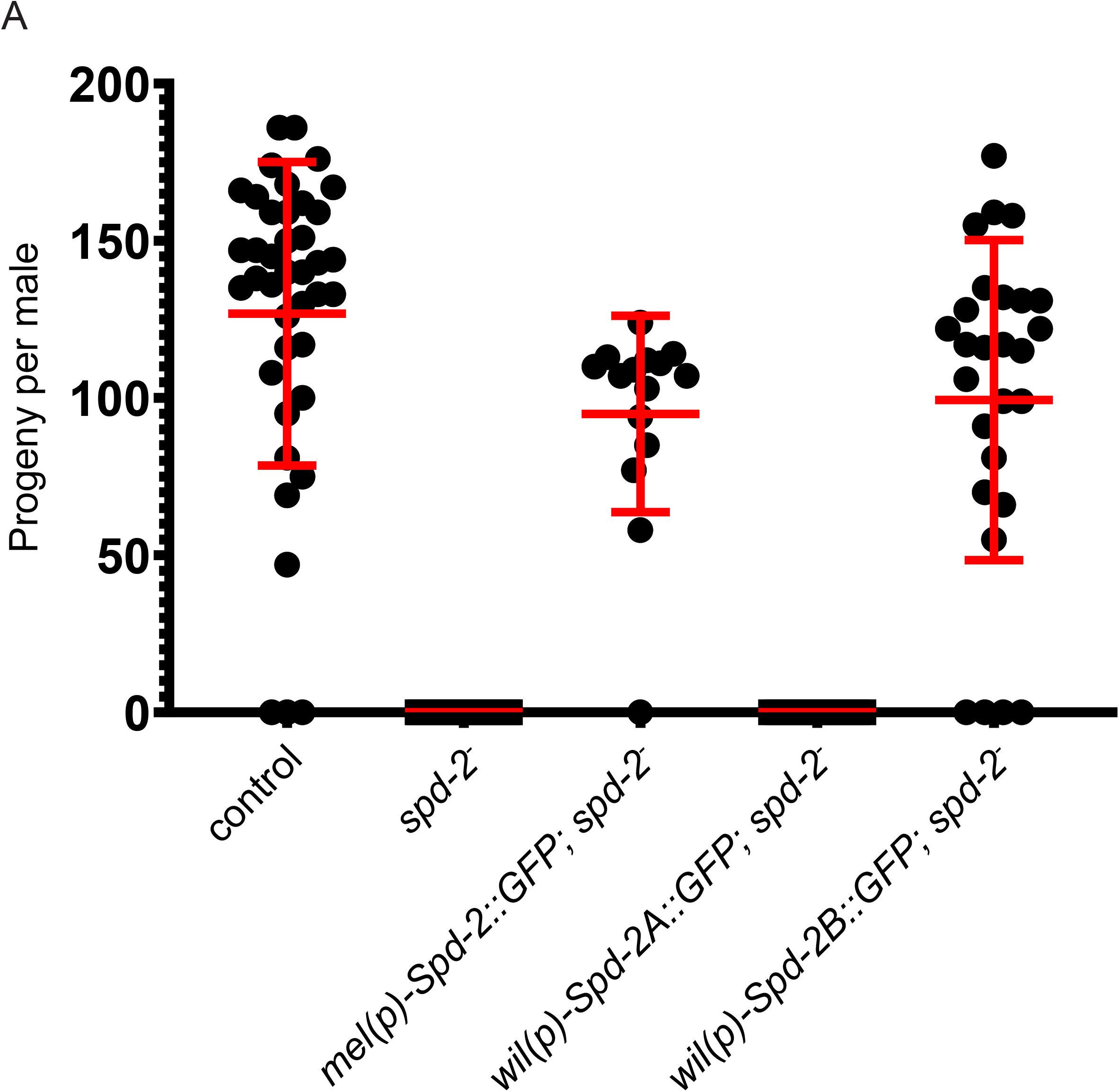
*Spd-2B* is required for male fertility but *Spd-2A* is dispensable. (A) Male fertility of control, *spd-2^-^* mutant, and native promoter rescues. Single males were mated to two virgin females, and the total number of offspring per male was counted. Controls, *mel(p)-Spd-2*::GFP rescues and *wil(p)-Spd-2B::GFP* rescues were fertile, whereas *spd-2^-^* mutants and *wil(p)-Spd-2A::GFP* rescues were completely sterile.

Next, we ectopically overexpressed *UAS* transgenes in wild type spermatocytes using *topi-GAL4* to increase protein levels during meiosis and potentially force centrosomal localization. Spd-2::GFP had robust centriolar localization in spermatocytes which expanded into the PCM during meiosis (Figure 8A). Spd-2A::GFP was weakly localized to centrioles and meiotic PCM of spermatocytes (Figure 8B). In contrast, Spd-2B::GFP had robust centriolar localization and expansion into meiotic PCM (Figure 8C). We also tested whether *topi-GAL4*-driven *UAS* transgenes could rescue *spd-2^-^* meiotic spindle organization. *topi-GAL4* driving *UAS-Spd-2::GFP* fully rescues spindles (Figure 8D, E). In contrast, *topi-GAL4* driving *UAS-Spd-2A::GFP* fails to rescue spindles (Figure 8F). Finally, *topi-GAL4* driving *UAS-Spd-2B::GFP* fully rescues spindles (Figure 8G). Thus, even when ectopically overexpressed in spermatocytes Spd-2A localizes to meiotic centrosomes at low levels, does not expand into the PCM, and does not organize meiotic spindles, suggesting that *Spd-2A* has accumulated coding sequence mutations through the course of its evolution that led to loss of ancestral *Spd-2* meiotic PCM functions.

**Figure 8.**
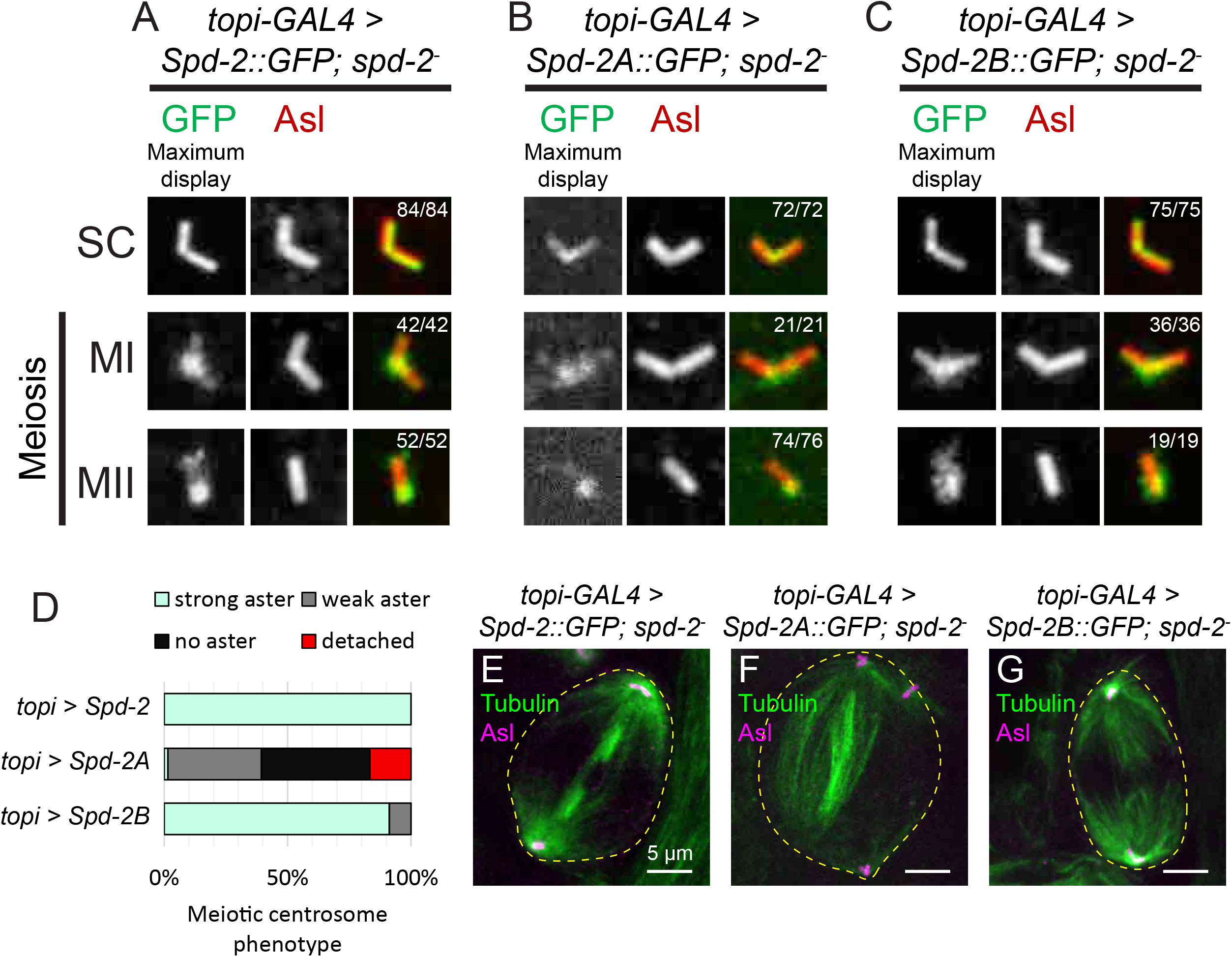
Spd-2A fails to properly localize to meiotic centrosomes. (A-C) Micrographs of *topi-GAL4* driving *UAS-Spd-2::GFP* (A), *UAS-Spd-2A::GFP* (B) and *UAS-Spd-2B::GFP* (C) showing Spd-2(A/B)::GFP localization to centrosomes in spermatocytes. GFP channels are displayed such that the dynamic range corresponds to the brightest and dimmest pixels. Note that both Spd-2 (A) and Spd-2B (C) are robustly recruited to spermatocyte centriole, whereas Spd-2A (B) is barely detectable. SC = spermatocytes; MI = meiosis I; MII = meiosis II. (D) Quantification of centrosome phenotypes for *topi-GAL4* rescues. Strong aster corresponds to centrosomes with full asters; weak asters correspond to centrosomes with only a few microtubules; no asters corresponds to centrosomes with no visible microtubules; and detached corresponds to centrosomes that have detached from the poles. (E-G) Micrographs of *spd-2^-^* mutants with *topi-GAL4* driving *UAS-Spd-2::GFP* (E), *UAS-Spd-2A::GFP* (F) and *UAS-Spd-2B::GFP* (G) stained for Tubulin (green) and Asl (magenta).

### Spd-2A C-terminal Tail Prevents Meiotic Spindle Organizing Function

To determine the specific amino acid changes responsible for the differences in Spd-2A and Spd-2B function we used chimeric transgenes as an initial attempt to map the meiotic function to a specific protein region. First, we combined the N-terminal half of Spd-2A with the C-terminal half of Spd-2B, and vice versa, to make *UAS-Spd-2A-B^chimera^* and *UAS-Spd-2B-A^chimera^*, respectively. *topi-GAL4* driving *UAS-Spd-2A-B^chimera^::GFP* fully rescues *spd-2^-^* meiotic spindle defects (Figure 9A, B), whereas *UAS-Spd-2B- A^chimera^::GFP* fails to rescue meiotic spindles (Figure 9C). This result suggests that the meiotic spindle-organizing function can be attributed to the C-terminal half of Spd-2B. The most conspicuous difference between Spd-2A and Spd-2B is the lack of the C-terminal-most ∼119 amino acids (the tail) in Spd-2B (Figure 2D). We therefore generated additional transgenes where we removed the tail from Spd-2A (*UAS-Spd-2A^ΔC-tail^::GFP*) or added the tail to Spd-2B (*UAS-Spd2B^+C-tail^::GFP*). We found that *topi-GAL4* driving *UAS-Spd-2A^ΔC-tail^::GFP* fully rescued *spd-2^-^* meiotic spindle defects (Figure 9D). In contrast, *topi-GAL4* driving *UAS-Spd2B^+C-tail^::GFP* fails to rescue *spd-2^-^* meiotic spindle defects (Figure 9E). Thus, the presence of the C-terminal tail of Spd-2A is responsible for its inability to organize meiotic PCM and spindles.

**Figure 9.**
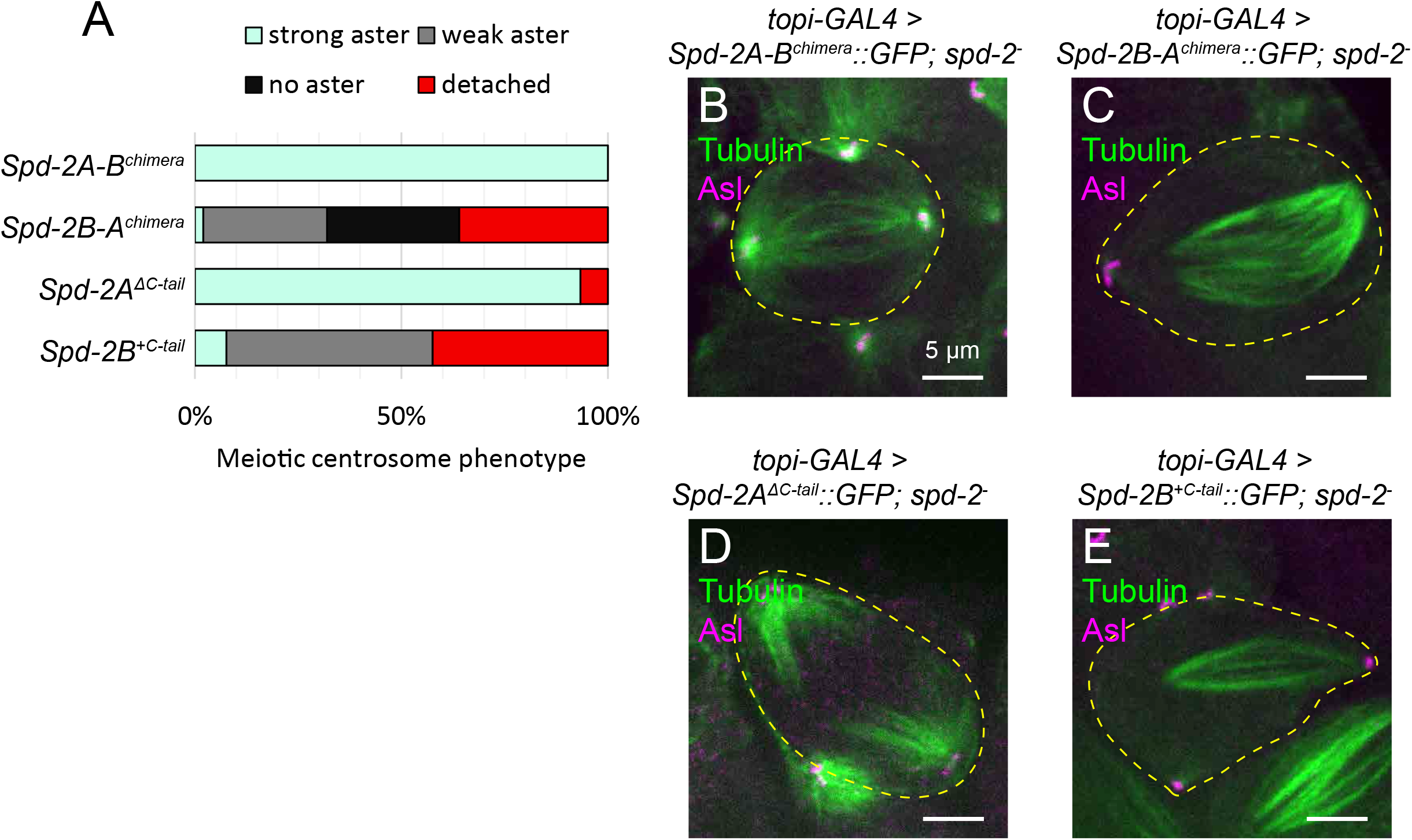
SApd-2A C-terminal tail inhibits meiotic spindle function. (A) Quantification of centrosome phenotypes for *topi-GAL4* > chimeric transgene rescues. Strong aster corresponds to centrosomes with full asters; weak asters correspond to centrosomes with only a few microtubules; no asters corresponds to centrosomes with no visible microtubules; and detached corresponds to centrosomes that have detached from the poles. (B-E) Micrographs of *spd-2^-^* mutants with *topi-GAL4* driving *UAS-Spd-2A-B^chimera^::GFP* (B), *UAS-Spd-2B-A^chimera^::GFP* (C), *UAS-Spd-2A^ΔC-tail^::GFP* (D) and *UAS-Spd-2B^+Spd-2A C-tail^::GFP* (E) stained for Tubulin (green) and Asl (magenta).

## Discussion

In this study, we sought to gain insight into the cell type-specific functions of centrosome proteins using an evolutionary approach. In *D. melanogaster*, *Spd-2* functions in both mitotic and meiotic divisions (Dix and Raff, 2007; Giansanti et al., 2008). The most counter-intuitive result from our study is that although Spd-2A is the more highly conserved paralog and contains all protein regions found in Spd-2, it has lost the ability to function in PCM recruitment in meiotic cells (Figure 6D). Instead, the less well-conserved Spd-2B is capable of expanding into the PCM in meiotic cells (Figure 6E), and in mitotic cells when ectopically expressed (Figure 4F). We also found that the C-terminal 119 amino acid tail of Spd-2A, which is absent from Spd-2B, is responsible for preventing meiotic function (Figure 9D, E), explaining why even higher, ectopic expression of Spd-2A is still unable to rescue meiotic function (Figure 8B). Interestingly, removing the tail from Spd-2A rescues meiotic function, indicating that the PCM recruitment capability of Spd-2A is not lost and must be located in another region of the protein. These results suggest a two-step process for ancestral (and *D. melanogaster*) Spd-2 in driving PCM assembly: Step 1) Spd-2 priming via tail autoinhibition relief; Step 2) Spd-2 activation for PCM recruitment (Figure 10A).

**Figure 10.**
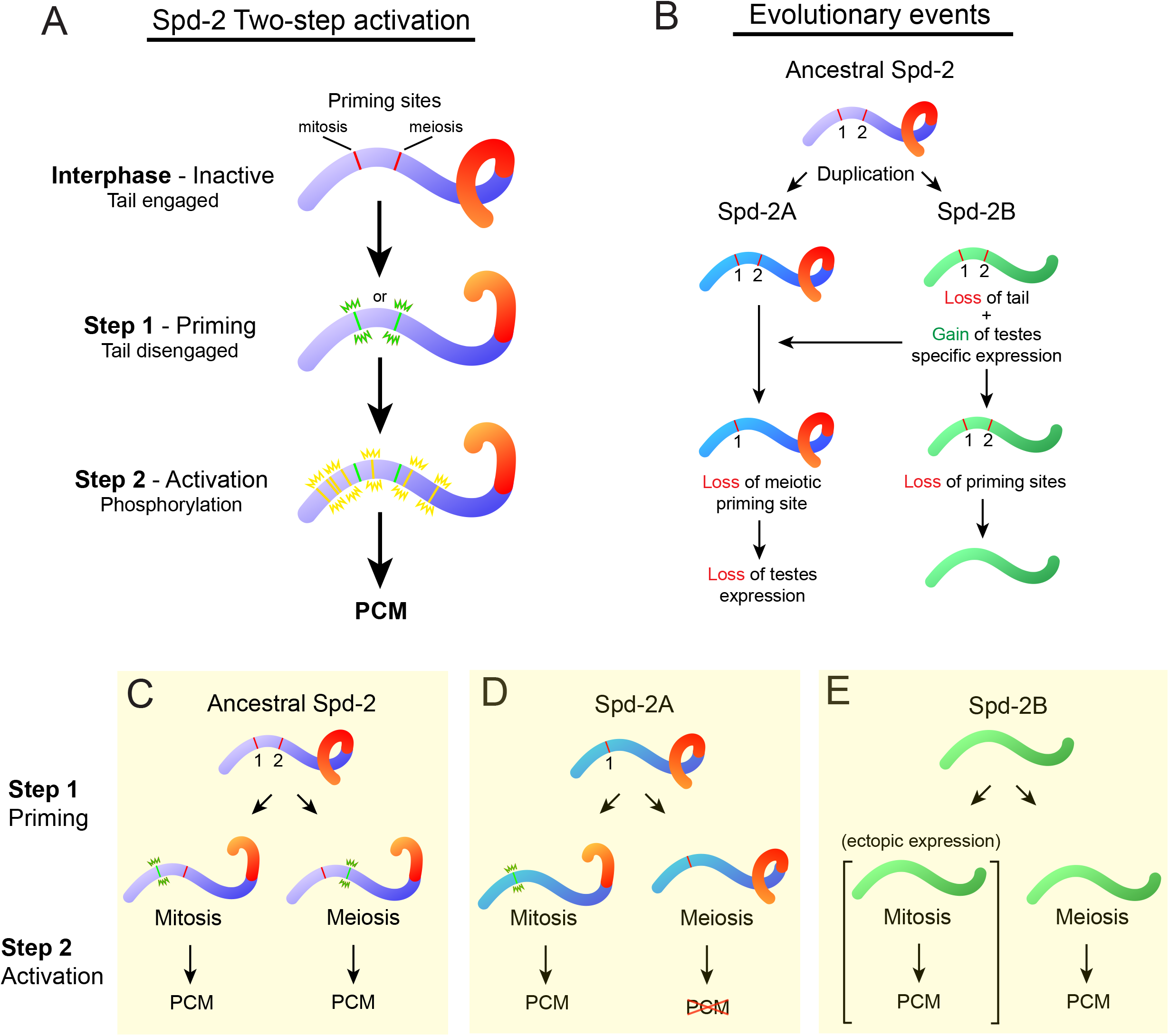
Hypothetical model for evolution of Spd-2A and Spd-2B cell type-specific specializations. (A) Hypothetical 2-step model for Spd-2 activation. In interphase, Spd-2 is unphosphorylated with the C-terminal tail preventing activation. In Step 1, Spd-2 is primed for activation by relieving the C-terminal tail autoinhibition and thus allowing phosphorylation. We speculate that this priming step is achieved via distinct mitotic and meiotic regulatory sites. In Step 2, Spd-2 is phosphorylated to allow recruitment of Polo and Cnn to expand the PCM. (B) Model of Spd-2 gene duplicate evolution. Ancestral Spd-2 is ubiquitously expressed and contains two distinct regulatory sites and the C-terminal tail. Gene duplication gave rise to Spd-2B, which gained testes-specific expression and lacked the C-terminal tail. Since Spd-2B was able to function in meiotic PCM assembly, there was no longer evolutionary pressure for Spd-2A to maintain the meiotic regulatory site or testes expression and thus they were lost. Similarly, we speculate that there would be no pressure to maintain either regulatory site in Spd-2B, and thus we predict that they would also be lost. (C) Ancestral Spd-2 is autoinhibited via the C-terminal tail during interphase. Priming is achieved via distinct regulatory sites during mitosis or meiosis. (D) Spd-2A, which retains the C-terminal tail, is properly primed during mitosis. However, due to loss of a hypothetical meiotic regulatory site, Spd-2A remains autoinhibited during meiosis, thus blocking phosphorylation and preventing PCM assembly. (E) Spd-2B does not require a priming step due to loss of the C-terminal tail. Spd-2B is not expressed in somatic cells and therefore does not normally contribute to mitotic PCM, however ectopic expression leads to proper PCM assembly. Spd-2B, which is expressed in male germline cells, can freely organize PCM during meiosis because it has no C-terminal tail to block phosphorylation.

Previous work on ‘Step 2’ in multiple systems showed that Spd-2/SPD-2/Cep192 are activated via phosphorylated during mitosis, and that this phosphorylation is required to efficiently recruit Polo/Plk1 and expand the mitotic PCM (Alvarez-Rodrigo et al., 2019; Decker et al., 2011; Meng et al., 2015). Our work does not shed light on ‘Step 2’ except to show that it does not require the C-terminal tail. Instead, our work has identified ‘Step 1’, a priming step that provides an additional level of Spd-2 regulation. Our results provide at least two plausible explanations for autoinhibition relief: A) autoinhibition by the tail is male meiosis-specific. In this case, Spd-2 priming is only necessary in meiosis, while in mitosis ‘Step 2’ is sufficient for PCM regulation. B) A second, more favorable explanation is that there are distinct cell type-specific (mitotic and meiotic) mechanisms of autoinhibition relief to prime Spd-2 (Figure 10A). The mechanism of relief could involve a cell type specific binding partner (a protein activator), or a cell type specific post-translational modification at a distinct mitotic residue(s) and meiotic residue(s) within Spd-2 (Figure 10A, priming sites). Data from the literature supports this two step model and provides many hypotheses that need future testing. For example, although evidence from *C. elegans* indicates that SPD-2 exists in a monomeric state in the cytoplasm, intramolecular Spd-2 interactions have been reported by several groups via yeast two-hybrid (Conduit et al., 2014; Galletta et al., 2016a). Thus, one possibility is that the priming step represents a switch from a monomeric state during interphase to an oligomeric state at the centrosome during mitosis, which somehow involves tail disengagement. Future work should aim to better understand the nature of this putative mechanism and why it appears to function differently across cell types.

*Spd-2B* likely arose via a small-scale, dispersed DNA-based duplication mechanism (Ezawa et al., 2011) as evidenced by the presence of introns and its location on a different chromosome from its parent gene. Testis expression is a common initial feature of new gene duplicates (Vinckenbosch et al., 2006), and new gene duplicates appear to frequently acquire regulatory elements that drive testis expression from neighbor genes (Bai et al., 2009). Interestingly, *D. melanogaster* high throughput expression data show that five of the seven genes in the *Spd-2B* syntenic region (Figure S3A) are most strongly expressed in the testes (Figure S3B-G), consistent with *Spd-2B* acquiring enhancers that drive male germline expression from its neighbor genes. Thus, the testis-specific expression pattern of *Spd-2B* and the testis-biased expression patterns of several neighbor genes suggests that *Spd-2B* picked up testis expression from neighbor gene regulatory elements or from the local chromatin environment at the moment of its birth (Figure 10B). Once Spd-2B had gained spermatocyte expression and lost the C-terminal tail, there would no longer be any evolutionary pressure to maintain the putative autoinhibitory mechanism in male meiosis, thus leading to its loss in Spd-2A (Figure 10B). In summary, our model suggests ancestral (and *D. melanogaster*) Spd-2 properly organizes PCM in mitosis and meiosis by priming via distinct regulatory sites (Figure 10C). In contrast, Spd-2A organizes PCM in mitosis but remains inactive during meiosis due to loss of the relevant regulatory site (Figure 10D). Finally, Spd-2B does not require a priming step due to loss of the C-terminal tail. Therefore, Spd-2B is capable of organizing PCM in both meiosis and mitosis, although under normal conditions it is only expressed in meiotic cells (Figure 10E).

Our discoveries with Spd-2A and Spd-2B suggest that mitotic and male meiotic cell division have subtly different requirements for PCM assembly. One major difference between mitotic and male meiotic cell division in *Drosophila* is the relatively large size of the meiotic spindle. The large size of the male meiotic spindle is thought to be due in part to high concentrations of tubulin monomers in the spermatocyte cytoplasm required for sperm tail axoneme formation in later spermatogenesis (Lattao et al., 2012). While somatic cells lacking centrosomes can still progress through mitosis using chromatin-nucleated microtubules, male germline cells lacking centrosomes fail to progress through meiosis (Gatti et al., 2012). This failure is thought to reflect a requirement for rapid centrosome-driven polymerization of microtubules in the tubulin-rich spermatocytes. Thus, one possibility is that, in *D. melanogaster*, Spd-2 activity must be upregulated during meiosis to enhance its ability to nucleate microtubules, whereas, in *D. willistoni*, Spd-2B is simply specialized for rapid microtubule nucleation to support this distinct requirement of male meiosis.

Our study also revealed several other centrosome gene duplications in the *Drosophila* lineage. *Cnn*, which encodes the other major PCM protein in *Drosophila*, underwent two independent duplications; learning whether these *Cnn* duplicates are specialized in a manner similar to *Spd-2A* and *Spd-2B* could further support different requirements for PCM in male meiosis. Additionally, of particular interest are duplications of *Asl* and *Cp110*, which were both duplicated multiple times in the *obscura* group, providing the potential for multiple specialized forms of these centrosome proteins. Overall, evolution has provided for us a rich resource for uncovering new insights into centrosome biology.

## Methods

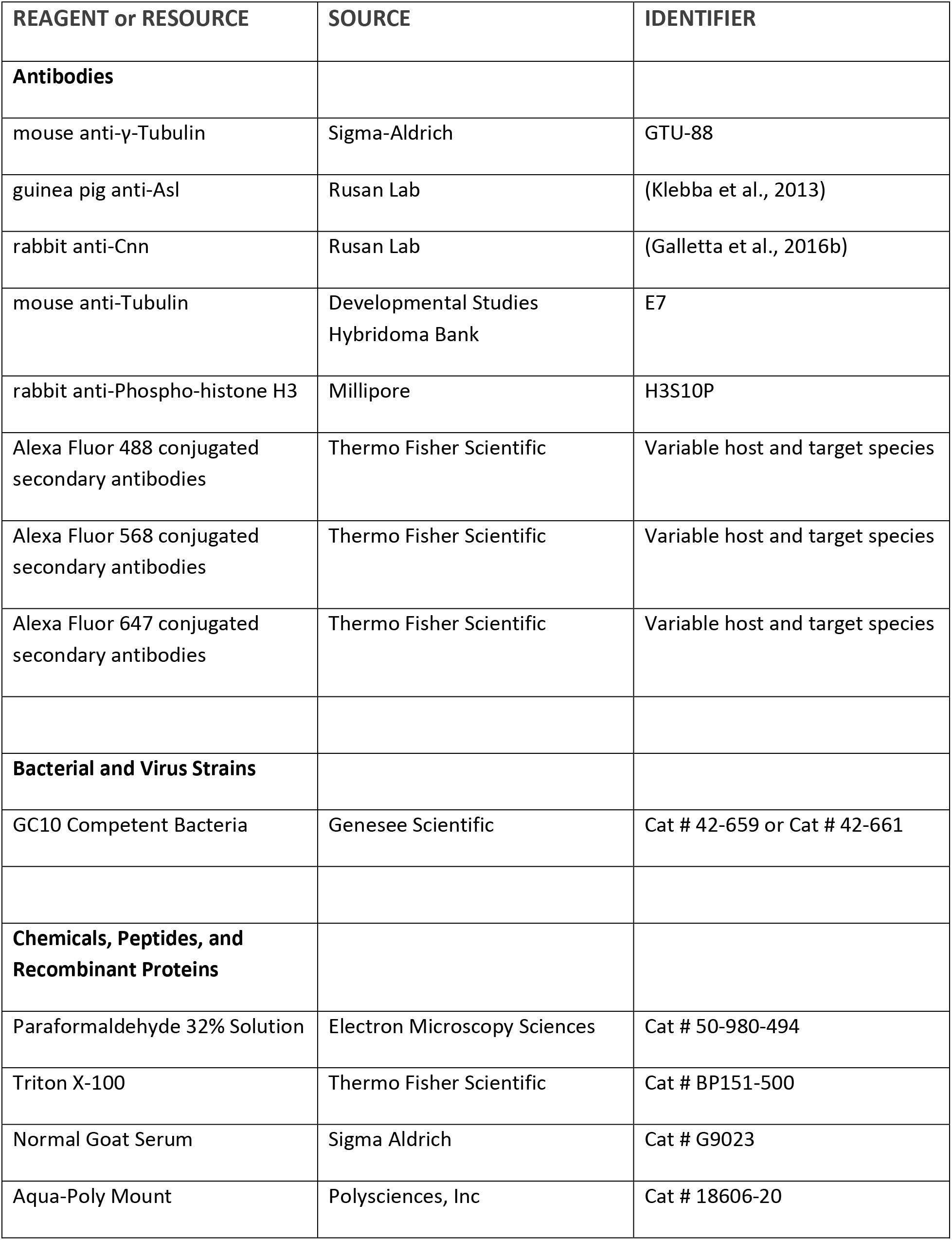
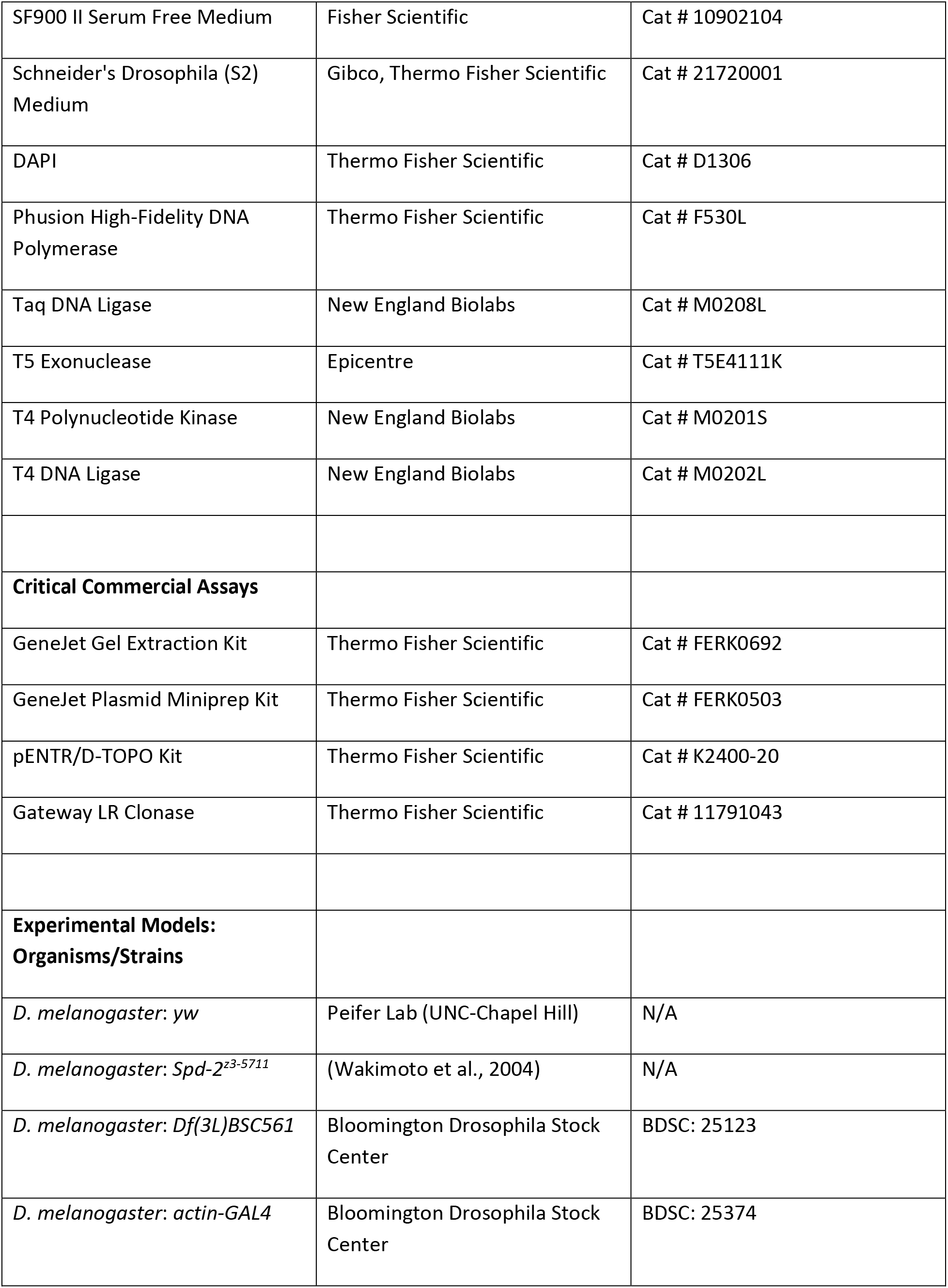
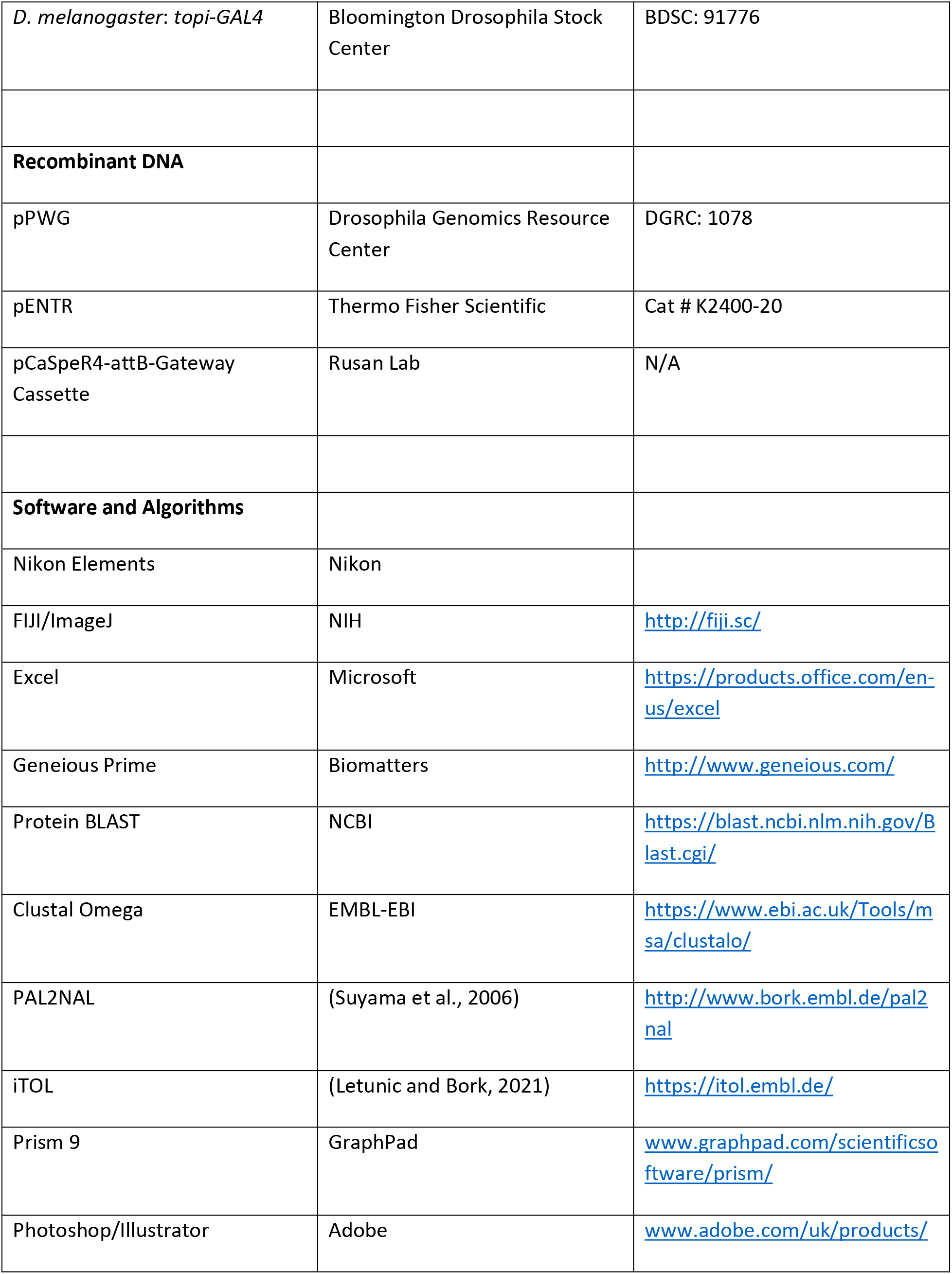
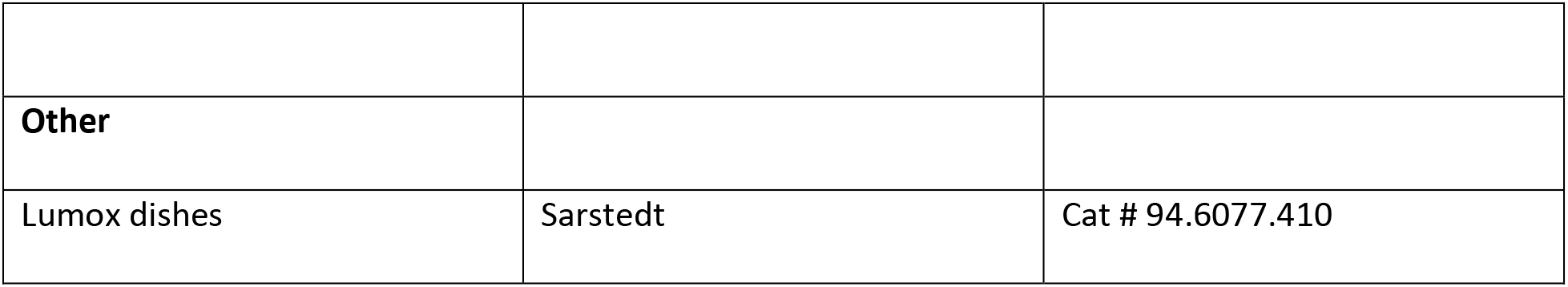

| REAGENT or RESOURCE | SOURCE | IDENTIFIER |
| --- | --- | --- |
| **Antibodies** |  |  |
| mouse anti-γ-Tubulin | Sigma-Aldrich | GTU-88 |
| guinea pig anti-Asl | Rusan Lab | (Klebba et al., 2013) |
| rabbit anti-Cnn | Rusan Lab | (Galletta et al., 2016b) |
| mouse anti-Tubulin | Developmental Studies Hybridoma Bank | E7 |
| rabbit anti-Phospho-histone H3 | Millipore | H3S10P |
| Alexa Fluor 488 conjugated secondary antibodies | Thermo Fisher Scientific | Variable host and target species |
| Alexa Fluor 568 conjugated secondary antibodies | Thermo Fisher Scientific | Variable host and target species |
| Alexa Fluor 647 conjugated secondary antibodies | Thermo Fisher Scientific | Variable host and target species |
| **Bacterial and Virus Strains** |  |  |
| GC10 Competent Bacteria | Genesee Scientific | Cat # 42-659 or Cat # 42-661 |
| **Chemicals, Peptides, and Recombinant Proteins** |  |  |
| Paraformaldehyde 32% Solution | Electron Microscopy Sciences | Cat # 50-980-494 |
| Triton X-100 | Thermo Fisher Scientific | Cat # BP151-500 |
| Normal Goat Serum | Sigma Aldrich | Cat # G9023 |
| Aqua-Poly Mount | Polysciences, Inc | Cat # 18606-20 |
| SF900 II Serum Free Medium | Fisher Scientific | Cat # 10902104 |
| Schneider's Drosophila (S2) Medium | Gibco, Thermo Fisher Scientific | Cat # 21720001 |
| DAPI | Thermo Fisher Scientific | Cat # D1306 |
| Phusion High-Fidelity DNA Polymerase | Thermo Fisher Scientific | Cat # F530L |
| Taq DNA Ligase | New England Biolabs | Cat # M0208L |
| T5 Exonuclease | Epicentre | Cat # T5E4111K |
| T4 Polynucleotide Kinase | New England Biolabs | Cat # M0201S |
| T4 DNA Ligase | New England Biolabs | Cat # M0202L |
| **Critical Commercial Assays** |  |  |
| GeneJet Gel Extraction Kit | Thermo Fisher Scientific | Cat # FERK0692 |
| GeneJet Plasmid Miniprep Kit | Thermo Fisher Scientific | Cat # FERK0503 |
| pENTR/D-TOPO Kit | Thermo Fisher Scientific | Cat # K2400-20 |
| Gateway LR Clonase | Thermo Fisher Scientific | Cat # 11791043 |
| **Experimental Models:**<br>**Organisms/Strains** |  |  |
| *D. melanogaster: yw* | Peifer Lab (UNC-Chapel Hill) | N/A |
| *D. melanogaster: Spd-2<sup>z3-5711</sup>* | (Wakimoto et al., 2004) | N/A |
| *D. melanogaster: Df(3L)BSC561* | Bloomington Drosophila Stock Center | BDSC: 25123 |
| *D. melanogaster: actin-GAL4* | Bloomington Drosophila Stock Center | BDSC: 25374 |
| *D. melanogaster: topi-GAL4* | Bloomington Drosophila Stock Center | BDSC: 91776 |
| **Recombinant DNA** |  |  |
| pPWG | Drosophila Genomics Resource Center | DGRC: 1078 |
| pENTR | Thermo Fisher Scientific | Cat # K2400-20 |
| pCaSpeR4-attB-Gateway Cassette | Rusan Lab | N/A |
| **Software and Algorithms** |  |  |
| Nikon Elements | Nikon |  |
| FIJI/ImageJ | NIH | http://fiji.sc/ |
| Excel | Microsoft | https://products.office.com/en-us/excel |
| Geneious Prime | Biomatters | http://www.geneious.com/ |
| Protein BLAST | NCBI | https://blast.ncbi.nlm.nih.gov/Blast.cgi/ |
| Clustal Omega | EMBL-EBI | https://www.ebi.ac.uk/Tools/msa/clustalo/ |
| PAL2NAL | (Suyama et al., 2006) | http://www.bork.embl.de/pal2nal |
| iTOL | (Letunic and Bork, 2021) | https://itol.embl.de/ |
| Prism 9 | GraphPad | www.graphpad.com/scientificsoftware/prism/ |
| Photoshop/Illustrator | Adobe | www.adobe.com/uk/products/ |
| **Other** |  |  |
| Lumox dishes | Sarstedt | Cat # 94.6077.410 |

### Lead Contact and Materials Availability

Further information and requests for reagents should be directed to and will be fulfilled by the Lead Contact, N.M.R. All unique/stable reagents generated in this study are available upon request without restriction.

### BLAST Screen for Gene Duplications

The protein sequences for 13 major centrosome proteins were obtained from FlyBase (Larkin et al., 2021). For genes with multiple splicing variants, the variant that included all exons was chosen for the BLAST screen (Supplemental File 1). Each protein sequence was then used as a query for Protein BLAST (McGinnis and Madden, 2004) against 35 sequenced *Drosophila* species (Supplemental File 1). Results were visually inspected, and positive hits were used in a reverse protein BLAST search to determine whether they returned the original query as the top hit. Detailed sequences and records of the screen can be found in Supplemental File 1.

To determine synteny, the gene region including 10 kb upstream and downstream of either Spd-2A or Spd-2B was obtained from FlyBase and used as query for nucleotide BLAST against the *D. melanogaster* genome.

### Sequence Analysis

The protein sequence and coding sequence for Spd-2 homologs in 7 species were obtained from FlyBase. Protein sequences were first aligned using Clustal Omega with default settings (Sievers et al., 2011). The protein alignment was then converted to a codon alignment using PAL2NAL (Suyama et al., 2006). Gaps were removed from the alignment, and Clustal Omega was used to generate a tree, which was visualized using iTOL (Letunic and Bork, 2021).

To compare both Spd-2A and Spd-2B to *D. melanogaster* Spd-2, the proteins were divided into an N-terminal half, C-terminal half, and a C-terminal tail that was absent from Spd-2B. Spd-2A and Spd-2B protein segments were aligned with the corresponding segment of Spd-2 using Clustal Omega, and the total number of identical amino acids and the total number of inserted or deleted amino acids was recorded.

### PCR Screen in *Willistoni* Group Species

*Drosophila paulistorum, Drosophila equinoxialis, Drosophila tropicalis, Drosophila nebulosa, Drosophila sucinea*, and *Drosophila virilis* strains were obtained from the Drosophila Species Stock Center (Cornell College of Agriculture and Life Science). Genomic DNA was extracted and used to amplify *Spd-2A* and *Spd-2B* from each species. For members of the *Willistoni* group, primers were based off of *D. willistoni* sequences. Sanger sequencing was used to confirm the identity of PCR products and rule out the possibility of DNA contamination from *D. willistoni*.

### D. melanogaster

Experimental fly crosses were maintained on Bloomington Recipe Fly food from LabExpress and kept at 25°C. Crosses were either 8 virgin females per vial or 20 virgin females per bottle, with at least half as many males. Controls were *yw* crossed to *Df(3L)BSC561*, and *spd-2^-^* were *Spd-2^z3-5711^* crossed to *Df(3L)BSC561*. All new transgenic animals were generated using standard embryo injection protocols by BestGene (Chino Hills, CA).

### Generation of Transgenic *Drosophila*

For native promoter *Spd-2*, *Spd-2A* and *Spd-2B* transgenes, the relevant gene region from the end of the upstream neighbor gene to the beginning of the downstream neighbor gene was PCR amplified from genomic DNA and cloned into pENTR. Gibson assembly was used to insert a GFP-myc or GFP-HA attB cassette in frame at the 3’ end of the coding sequence. Genes were moved to pCaSpeR4 containing an attB recombination site and a Gateway cassette by LR Clonase reactions. pCaSpeR4-attB-Spd-2 constructs were then injected into *D. melanogaster* strain *y^1^ v^1^; P[CaryP]attP40* which contains an insulated landing site.

For UAS-Spd-2::GFP, UAS-Spd-2A::GFP, and UAS-Spd-2B::GFP, the relevant coding sequences (including introns) were PCR amplified from genomic DNA and cloned into pENTR using standard Gateway cloning methods. Gateway cloning was used to move constructs from pENTR into pPWG. These constructs were then injected into *D. melanogaster* strain *yw*.

### Immunostaining

For larval brain experiments, 3^rd^ instar larvae were used. For testes experiments, pupae of various ages were used. Tissues were dissected in SF900 S2 cell media and fixed with 8% paraformaldehyde in SF900 for 20 minutes at room temperature (RT). Fixed samples were washed three times in PBST (PBS + 0.2% Triton-X100) and blocked for at least 1 hour in PBST + 5% normal goat serum (NGS). Blocked samples were incubated in PBST + NGS with primary antibody for either 2 hours at RT or else overnight at 4°C. Samples were washed five times in PBST + NGS, and then incubated in PBST + NGS with secondary antibody and DAPI for either 2 hours at RT or else overnight at 4°C. Samples were washed two times in PBST + NGS, two times in PBST, and then several washes in PBS to remove all Triton-X100. Samples were mounted on #1.5 coverslips and mounted in AquaPoly Mount.

### Light Microscopy and Image Analysis

Imaging was performed using a Nikon W1 spinning disc confocal equipped with a Prime BSI cMOS camera (Photometrics) and a 100X/1.4 NA silicon immersion objective, controlled using Nikon Elements software. Fluorescence intensities were measured and images were prepared for display using ImageJ software (Schneider et al., 2012).

### Male Fertility Assay

Bottles of *yw* with 10 females and 10 males were flipped daily to prevent over-crowding. Virgin *yw* females were collected from these bottles and aged for two days. Males of various genotypes to be tested were similarly collected and aged for two days, before being individually placed into vials with two virgin females to mate over the course of four days. After four days, the male was discarded, and females were allowed to lay eggs. At day 17, the total number of offspring per male was counted.

### Statistical analysis

Data analysis was performed using Microsoft Excel and GraphPad Prism. In all graphs, the mean ± standard deviation is presented. Shapiro-Wilk test was used to test the assumption of normality. ANOVA was used to determine significance, which is reported in Figure legends.

## Acknowledgements

We thank Bloomington Drosophila Stock Center and the National Drosophila Species Stock Center at Cornell College of Agricultural Sciences for fly stocks, and Developmental Studies Hybridoma Bank for antibodies. Carey Fagerstrom performed all cloning. We thank Alex Kelly, Takashi Akera, Matthew Hannaford, Ramya Varadarajan, and Chaitali Khan for helpful discussion.

---

**Figure S1.**
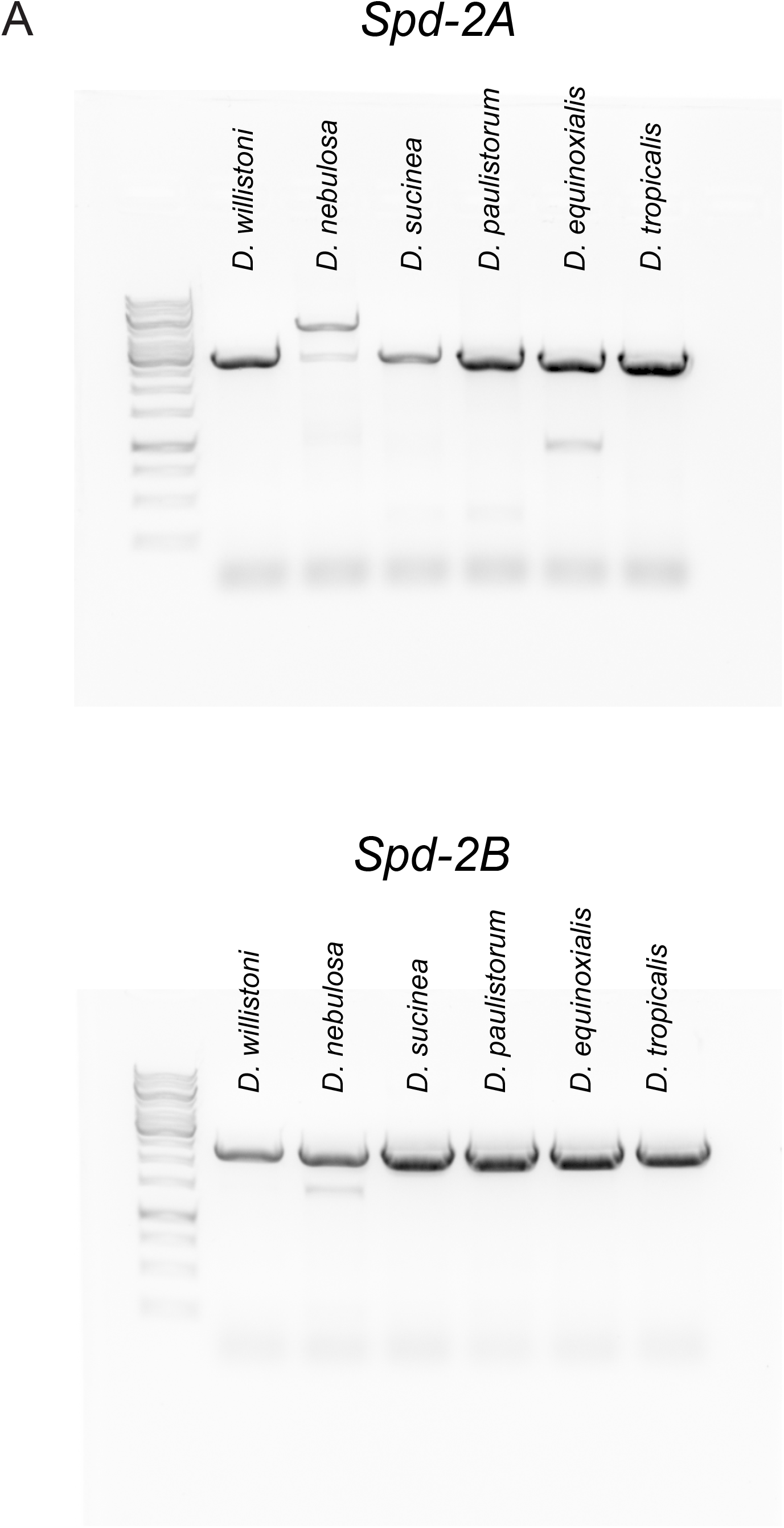
*Willistoni* group *Spd-2A* and *Spd-2B* PCR products. (A) Ethidium bromide-stained agarose gel showing PCR products for *Spd-2A* (top) and *Spd-2B* (bottom) from six *Willistoni* group species.

**Figure S2.**
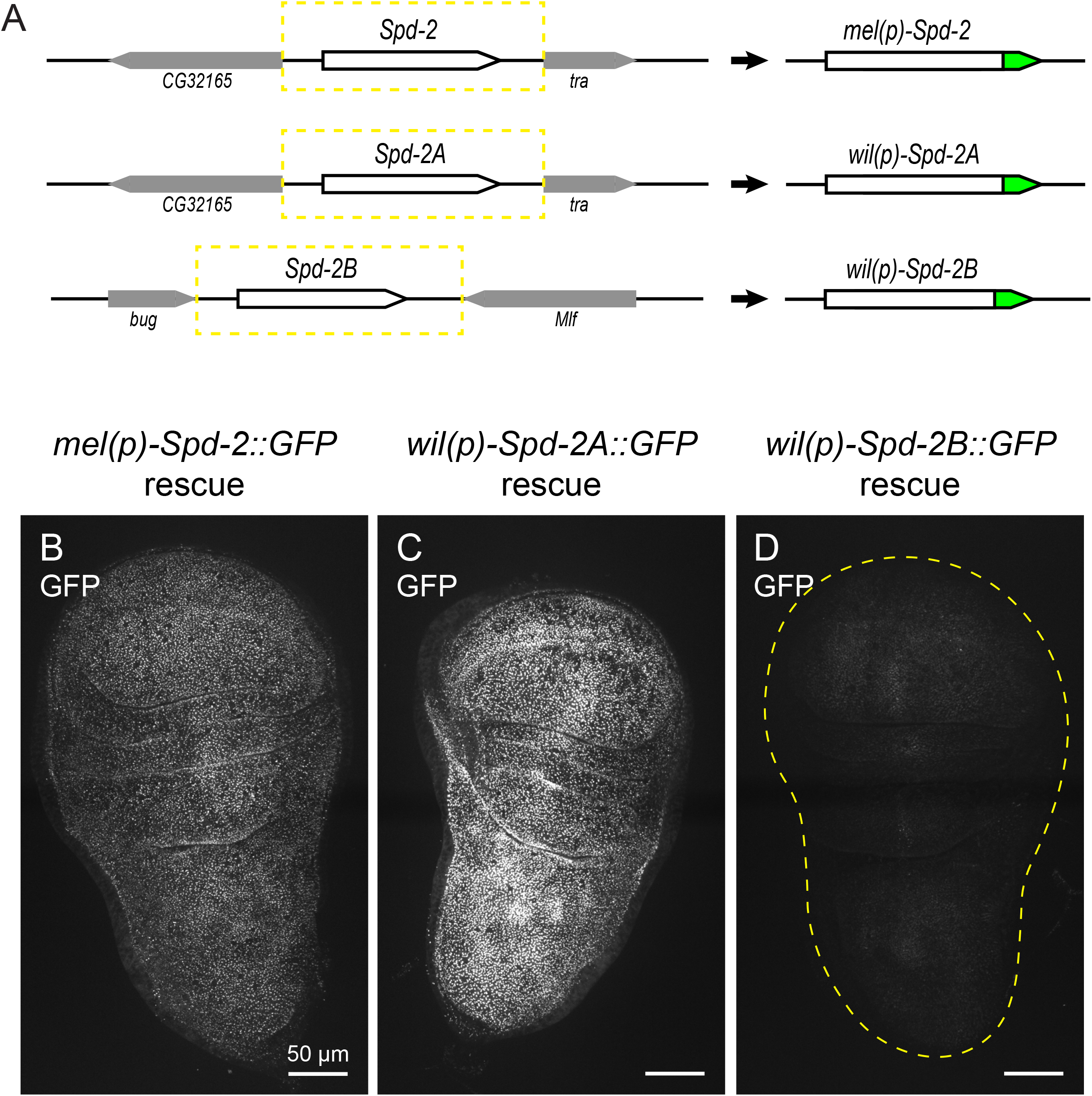
*Spd-2* and *Spd-2A*, but not *Spd-2B*, are expressed in somatic tissue. (A) Diagram of native promoter transgene design. Gene regions including upstream and downstream regulatory DNA for *D. melanogaster Spd-2* and *D. willistoni Spd-2A* and *Spd-2B* were used to generate GFP-tagged transgenes inserted into the attp40 landing site. (B-D) Micrographs of unfixed 3^rd^ instar larval wing discs expressing *mel(p)-Spd-2::GFP* (B), *wil(p)-Spd-2A::GFP* (C) and *wil(p)-Spd-2B::GFP* (D). Both Spd-2::GFP and Spd-2A::GFP are expressed in wing discs, whereas Spd-2B::GFP is undetectable.

**Figure S3.**
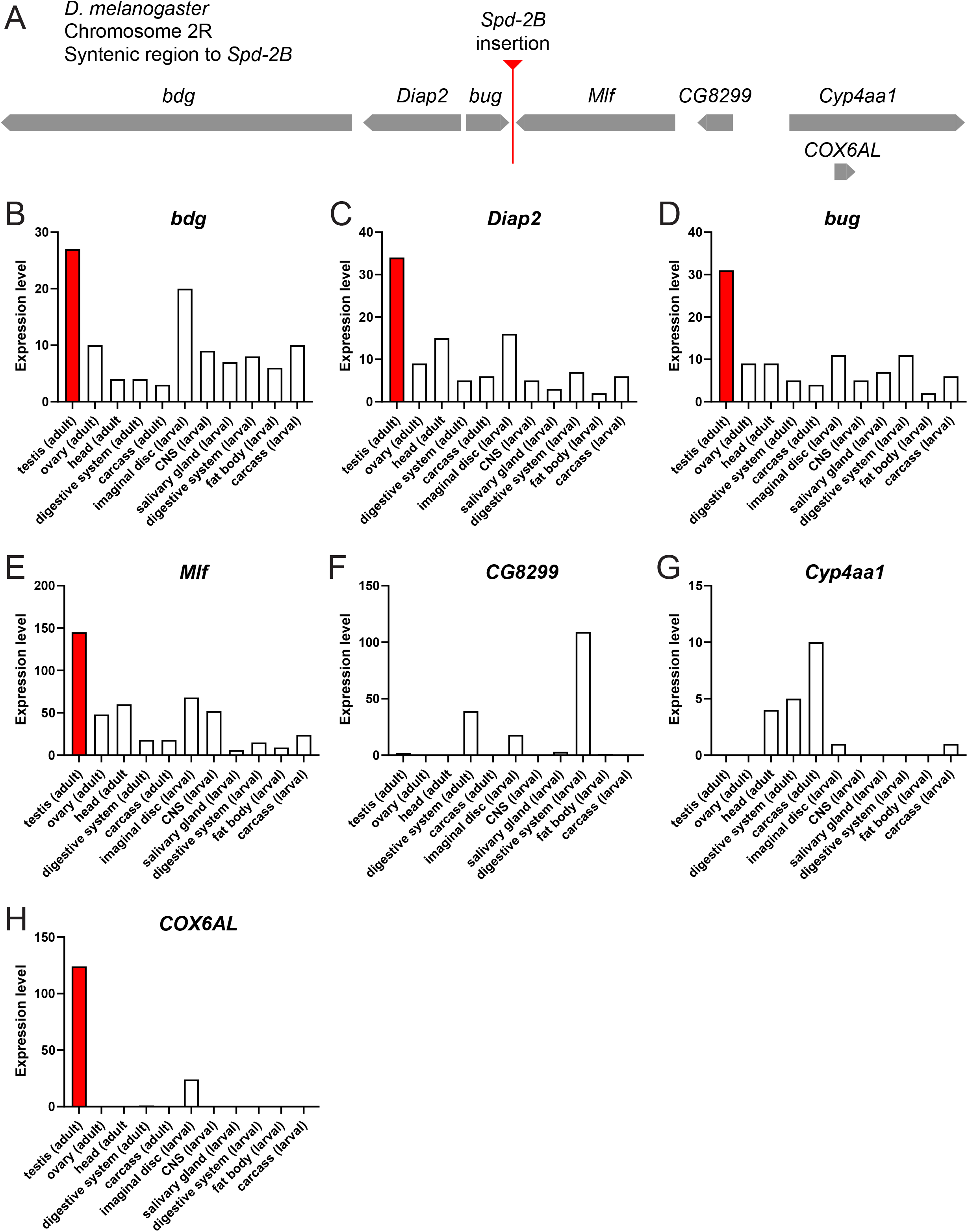
*Spd-2B* landed in a locus with testis-biased expression. (A) Representation of the *D. melanogaster* genome region that is syntenic to the locus where *Spd-2B* was inserted. (B-H) Expression profiles of genes in the *Spd-2B* syntenic locus, showing testis expression in red. Five out of seven genes near the Spd-2B insertion site have testis-biased expression. Expression levels were obtained from Flybase high-throughput tissue-specific expression data.

